# Twist - torsion coupling in beating axonemes

**DOI:** 10.1101/2024.03.18.585533

**Authors:** Martin Striegler, Benjamin M. Friedrich, Stefan Diez, Veikko F. Geyer

## Abstract

Motile cilia and flagella are ubiquitous cell appendages whose regular bending waves pump fluids across tissue surfaces and enable single-cell navigation. Key to these functions are their non-planar waveforms with characteristic torsion. It is not known how torsion, a purely geometric property of the shape, is related to mechanical deformations of the axoneme, the conserved cytoskeletal core of cilia and flagella. Here, we assess torsion and twist in reactivated axonemes isolated from the green alga *Chlamydomonas reinhardtii*. Using defocused darkfield microscopy and beat-cycle averaging, we resolve the 3D shapes of the axonemal waveform with nanometer precision at millisecond timescales. Our measurements reveal regular hetero-chiral torsion waves propagating base to tip with a peak-to-peak amplitude of 22 º/µm. To investigate if the observed torsion results from axonemal twist, we attach gold nanoparticles to axonemes to measure its cross-section rotation during beating. We find that locally, the axonemal cross-section co-rotates with the bending plane. This co-rotation presents the first experimental evidence for twist-torsion coupling and indicates that twist waves propagate along the axoneme during beating. Our work thus links shape to mechanical deformation of beating axonemes, informing models of motor regulation that shape the beat of motile cilia.

## Introduction

Cilia are slender, membrane-enclosed protrusions of eukaryotic cells. Motile cilia drive essential physiological functions such as the breaking of the left-right symmetry in mammalian embryos ^1^, fluid transport in the human respiratory tract, fallopian tubes, and cerebral cavities ^2–4^ as well as the locomotion of single cell micro swimmers like sperm and algae. The internal mechanical core of cilia is evolutionary conserved among eukaryotes and is called the axoneme. The axoneme consists of nine doublet microtubules (DMTs), which are cylindrically arranged around a pair of singlet microtubules (which scaffold the central apparatus) (**Figure 1A**). Dynein motors are distributed along the entire length of the axoneme and arranged in a chiral fashion, and, when active, slide adjacent DMTs. This sliding is constrained at the ciliary base and along the axoneme, which converts sliding into traveling waves of bending that shape the ciliary waveform.

**Figure 1:**
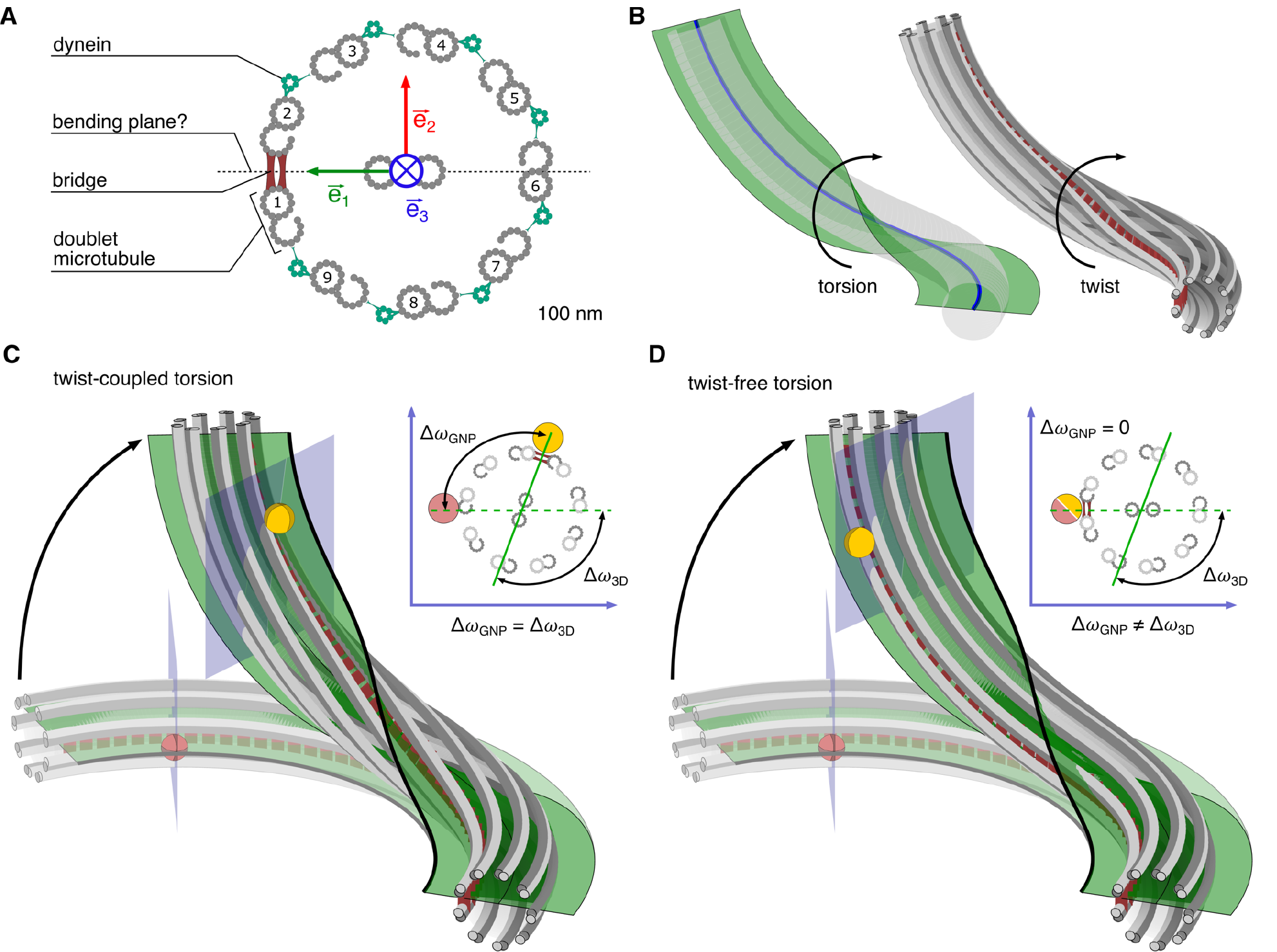
Twist and Torsion in axonemes. **(A)** Schematic of axoneme cross-section (viewed from base) with numbered DMTs 1-9 ^14^. The bridge (red) is between DMT1 and DMT2. According to the “rigid-bridge-hypothesis”, the bridge sets the bending plane. The green and red arrows indicate the material frame of the axoneme (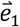 and 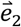 span the cross-sectional plane, with 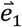 pointing towards the bridge and 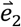 being orthogonal to 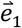, while 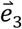 is normal to the cross-section, pointing along the centerline). Dynein motors (green) are permanently attached to one DMT and transiently interact with their clockwise neighbor. **(B)** Cartoon of a twisted axoneme in which torsion and twist are coupled. Left panel: torsion quantifies the rotation of the bending plane (green plane) along the 3D centerline (blue line). Right panel: twist quantifies the rotation of the cross-section in the material frame (cross-section orientation is marked by a dashed red line, indicating the bridge). **(C)** Expected movement of an axoneme-attached gold nanoparticles (GNP) assuming the “rigid-bridge-hypothesis” is true. In this scenario, the rotation of the bending plane (green plane with black outline) by an angle Δω_3D_ is exclusively due to axonemal twist. Note: the helical shape of the dashed red line (bridge) shows that the cross-section (with attached GNP) rotates by the same amount as the bending plane (Δ*ω*_GNP_ = Δ*ω*_3D_). The GNP-plane in which the rotation of the local cross-section *ω*_GNP_ and the rotation of the bending plane *ω*_3D_ is measured (as seen from the laboratory coordinate system), lies normal to the xy plane. **(D)** Alternative scenario of twist-free torsion of the axoneme, where the rotation of the bending plane (green plane with black outline) is caused by bending into different directions relative to the bridge (dashed red line). Note: the dashed red line (bridge) shows no cross-section rotation. In this scenario, the rotation of the cross-section (with attached GNP) by an angle Δ*ω*_GNP_ is different from the rotation angle of the bending plane Δ*ω*_3D_. The transparent axoneme (light gray) in C and D depict a planar reference shape (no torsion) with the reference position of the GNP. The insets in panels C and D show schematic of axonemal cross-sections in the GNP plane. The red and yellow spheres show the GNP in its reference position (planar shape) and its final position (non-planar shape) respectively.

Ciliary waveforms are often non-planar. This non-planarity supports their physiological functions. Theoretical studies have shown that non-planarity improves fluid pumping ^5–7^. In single cells, non-planar beat patterns cause cell rotation and helical swimming, which is key to chemo- and phototactic navigation strategies along chiral paths ^8–11^ as well as rheotaxis ^10,12,13^. How the non-planarity in ciliary waveforms is generated and how it relates to internal deformations of the axonemal structure is not known.

To investigate the relation between non-planarity and structural deformations in axonemal shapes, the concepts of twist and torsion need to be carefully distinguished: (i) Twist describes the internal rotation of the axoneme along the arc-length and characterizes the structural deformation of the axoneme’s material and (ii) Torsion describes the rotation of the bending plane along the arc-length and is a mathematical property of 3D curves. When twist and torsion are strictly coupled, twist causes an equal amount of torsion for a bent object (**Figure 1B**) and both terms are equivalent. This, for example, is the case for an object like an ordinary ruler, that has a highly anisotropic bending stiffness which sets a preferred bending plane (e.g. it only bends in one, but not the other direction). The axoneme, however, is a filament bundle where the degree of anisotropy in bending stiffnesses is unknown and thus this coupling is non-trivial.

The popular “rigid-bridge hypothesis” asserts that the bridge, an internal axonemal component which links DMT1 and DMT2, prevents sliding between these DMTs ^14^. If this hypothesis were true, any rotation of the bending plane of the axoneme (i.e. torsion) would necessarily result from an internal rotation of the axoneme, i.e., from twist (**Figure 1C**). On the contrary, if the sliding between DMT1 and DMT2 were fully unconstrained, the bending plane would be free to rotate. This would allow for a scenario where the axoneme assumes a non-planar shape but does not twist (**Figure 1D**). Thus, the question arises if torsion and twist are coupled, and if the rigid-bridge hypothesis is true.

To test for twist-torsion coupling, both quantities need to be measured in the same system and compared to each other. Whereas several reports indicate torsion in beating cilia and axonemes ^8,9,15–22^, twist cannot be assessed in a straight forward way. Note, torsion measurements alone cannot prove the existence of twist without access to the internal structure of the axoneme.

To date little is known about the twist in beating axonemes. Previous accounts for axonemal twist mostly came from static samples. For example, electron microscopic measurements on *Paramecium* cilia revealed heterochiral twist of 15-20 º/µm ^23^. The only dynamic measurement to date is an observation by ^24^, where a mitochondrion attached to a beating quail sperm tail served as a tracer particle to follow rotations of the axoneme. However, neither of these experiments quantified torsion and thus cannot test twist-torsion coupling.

To investigate if twist and torsion are coupled, we use reactivated axonemes purified from the green alga *Chlamydomonas reinhardtii* as a model. We measure the 3D waveforms with defocused-darkfield-microscopy and use a novel beat-cycle averaging method to achieve high spatio-temporal precision. This allows us to measure torsion reliably. Furthermore, we pioneer a rigorous error estimate, providing a space-time map of axonemal torsion inside a region-of-trust. To measure local rotations of the axonemal cross-section, we attach gold nanoparticles (GNP) as tracers to the outside of axonemes. By comparing these local cross-section rotations to the local rotations of the bending plane, we show that twist and torsion are coupled. This result relates the geometry of the 3D centerline of the axoneme to its internal structural deformations. Our measurements are consistent with a wave of twist that travels base-to-tip along the axoneme. In future studies, these twist dynamics can be used to test models that predict the motor activity in the beating axoneme.

## Results

### High-precision average 3D waveform of isolated axonemes

We measured the three-dimensional (3D) shapes of reactivated axonemes (**Figure 2B**) isolated from *Chlamydomonas reinhardtii* cells (**Figure 2A**), with high temporal and spatial resolution using defocused high-speed darkfield microscopy ^16,25^. First, we used a filament tracking software ^26^ to determine the axonemal centerline in two dimensions (2D), characterized as *x, y* coordinates along the arc-length. To compute the *z*-coordinate of the axonemal centerline at each arc-length position, we exploited that in darkfield microscopy the apparent width of objects increases as they become defocused. Specifically, we determined the full-width-at-half-maximum (FWHM) of the darkfield-signal of axonemes (measured perpendicular to the centerline, **Figure 2B** and **D**) at each arc-length position and used the relation between the FWHM and the *z*-coordinate as a calibration curve (determined from axonemes immobilized to a glass coverslip, see **Figure 2C** and **E** and **Figure S1** for more detail). Using this calibration curve, we reconstructed the 3D shapes (**Figure 2F**) and obtained 3D waveforms, which comprise the periodic sequence of axonemal shapes. The tracking errors in the *x, y* and *z* coordinates were *σ*_*x,y*_ ≈ 7.3 nm and *σ*_*z*_ ≈ 33.8 nm, respectively (**Figure S1**).

**Figure 2:**
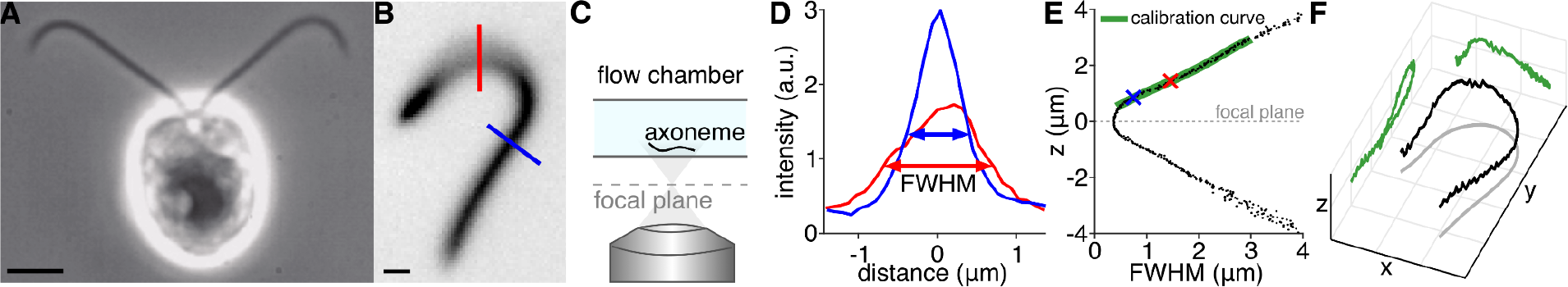
Reconstruction of 3D shapes from defocused high-speed darkfield microscopy images. **(A)** *Chlamydomonas reinhardtii* cell imaged with phase-contrast microscopy (scale bar 3 μm). **(B)** Defocused darkfield microscopy snap-shot of a reactivated axoneme with an exposure time of 1 ms (scale bar 1 µm). Colored lines indicate two positions where the axoneme is defocused (red) and where the axoneme is close to the focal plane (blue). **(C)** Schematic of defocused imaging. Images are recorded by focusing (focal plane is shown as the gray dashed line) below the axoneme sample. **(D)** Measurement of the full-width-at-half-maximum (FWHM) in intensity profiles along the colored lines in B, with lower FWHM values for axoneme parts closer to the focal plane. **(E)** Calibration of the relationship between FWHM and distance to focal plane. Values were obtained through *z*-scans of axonemes immobilized to the chamber surface closest to the objective (positive z-values correspond to axonemal positions above the focal plane). The calibration curve (green line) was obtained by a smoothing spline fit in MATLAB (smoothing parameter = 0.9). The red and blue crosses correspond to the curves and lines in panel D and B. **(F)** Example axoneme shape in 3D, with *z* coordinate obtained using the calibration curve in E.

Waveforms from individual axonemes were highly reproducible and thus allow to further increase the precision by averaging. We therefore defined a beat-cycle phase *ϕ* for each axonemal shape by fitting a periodic model function to each 2D shape (see Materials and Methods and **Figure S2A-D**). This method is superior to Fourier averaging because it is robust against frequency jitter. We then averaged the 3D shapes of 17 axonemes (with a total of 3755 beat-cycles and 14 frames/beat-cycle on average) with similar phase *ϕ*, so that we obtained a highly precise average waveform with increased temporal resolution (32 shapes/beat-cycle) (**Figure 3A, B**). The positional uncertainty of shapes in this average 3D waveform was 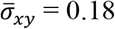 nm and 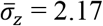 nm (standard error of mean; see **Figure S2K**). This is, to our knowledge, the measurement with the highest spatio-temporal precision of a beating axoneme or cilium to date.

**Figure 3:**
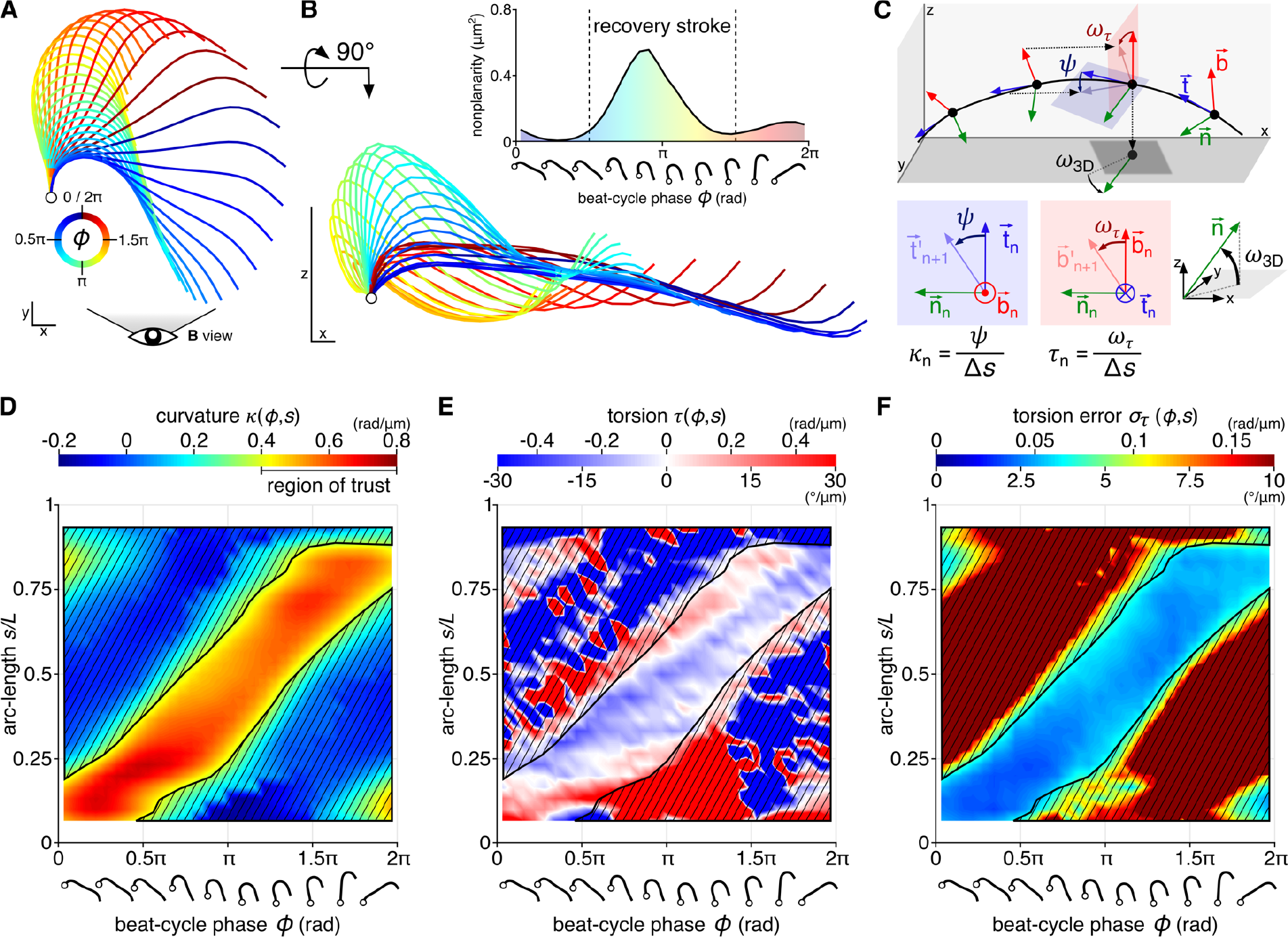
High-precision average 3D waveform of isolated axonemes and measurement of dynamic torsion. **(A)** *xy*-projection of average 3D waveform (average over 17 reactivated axonemes with total of 3755 beat-cycles), aligned at basal position. The color wheel represents the beat-cycle coordinate *ϕ* of each shape. Scale bar in *x* and *y* is 1 μm. **(B)** Side view of the average 3D waveform (*xz*-projection, panel A rotated by 90 º around the *x*-axis, scale bar is 1 μm, *z*-axis is normal to the boundary plane and points into the flow cell). Inset: nonplanarity of the waveform computed as the sum of the squared, normal residuals between each shape of the average waveform and a fitted plane. Scale bars in *x* and *z* are 500 nm. **(C)** Computation of torsion *τ*_n_ and 3D curvature *к*_n_ from the Frenet-Serret-frame with binormal vector 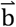 (red), normal vector 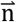 (green), and tangent vector 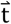 (blue) at subsequent arc-length positions (black filled circles) enumerated by *n* along the 3D centerline (black line), using the rotation angle of the local bending plane *ω*_*τ*_ and the in-plane rotation angle ψ of the tangent. Additionally, we measure *ω*_3*D*_ as the orientation of the normal vector with respect to the *xy*-plane of the laboratory frame. **(D)** Map of 3D curvature *к* as function of beat-cycle phase *ϕ* and arc-length *s*. **(E)** Map of torsion *τ* as function of beat-cycle phase *ϕ* and arc-length *s* (red shows dextral and blue sinistral torsion). **(F)** Map of estimated error *σ*_*τ*_ of torsion as function of beat-cycle phase *ϕ* and arc-length *s* (calculated using bootstrapping, for details see Figure S3). In panels D-F, hatched regions indicate where the curvature is below 0.4 rad/μm. The complementary, non-hatched region defines a region-of-trust for estimated torsion.

### Nonplanarity and torsion of the average 3D waveform

The average 3D waveform shows the strongest nonplanarity during the recovery stroke (**Figure 3B**, 0.5*π* < *ϕ* < 1.5*π*), which we quantified using a phase-dependent nonplanarity measure (**Figure 3B**, inset). The nonplanar centerline can also be quantified in terms of curvature and torsion. To do this, we use the Frenet-Serret frame, which is defined by three orthonormal vectors at each point along the arc-length: the tangent vector 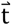, the normal vector 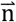 and the binormal vector 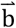 (**Figure 3C**). The 3D curvature *к* equals *ψ*/Δs, where *ψ* is the angle between subsequent tangent vectors 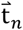 separated by an arc-length increment Δs. The torsion *τ* equals *ω*_*τ*_/Δs, where *ω*_*τ*_ is the angle between subsequent binormal vectors 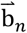 separated by an arc-length increment Δ s. Non-zero torsion indicates the rotation of the bending plane along the length (see also **Figure 1**). With this definition, *к* and *τ* have well-defined signs (up to a global choice of reference, see Materials and Methods).

Mathematically, torsion *τ* is only defined at arc-length positions where the curvature *κ* is non-zero. In practical terms, this implies that torsion estimates become unreliable when the curvature has low values. To identify regions, where we can reliably determine torsion, we locally quantify the variability of torsion estimates as function of absolute curvature |*κ*| (**Figure S3E**). We define a “region-of-trust” for reliable torsion estimates for curvature values above |*κ*| > 0.4 rad/μm where the estimated torsion error *σ*_*τ*_ is 4.2 °/μm or less (**Figure 3D-F**, non-hatched region in (*ϕ, s*)-space; more details see **Figure S3F**). From hereon, we only consider torsion values within this region-of-trust. Torsion can either be negative (blue: sinistral, i.e. left-handed) or positive (red: dextral, i.e. right-handed) and ranges from -19 °/μm to +21 °/μm. At a given arc-length position, torsion changes dynamically and occasionally even switches sign during one beat-cycle (**Figures 3E, S3G-I**). This becomes especially apparent at arc-length position *s* = 0.25 *L* or 0.75 *L*.

To characterize dynamic torsion, we estimate the peak-to-peak amplitude of the phase-dependent torsion for different arc-length positions. We find that this amplitude is approximately constant along arc-length with an average of 21.9 ± 5.8 °/μm (mean ± SD, N = 24, **Figure S4B**). Although this value only provides a lower bound, as we exclusively consider torsion within the region-of-trust, it shows that torsion changes with phase, i.e. that torsion is dynamic. For a given beat-cycle phase, torsion also changes as function of arc-length (**Figures 3E, S3J-L**). For example, at *ϕ* = 0.5 *π* and *ϕ* = *π* the sign of torsion changes twice along the arc-length. Generally, adjacent regions in (*ϕ, s*)-space of torsion with different signs are separated by bands of zero torsion (white regions in **Figure 3E**). The shape and orientation of these bands are reminiscent of a traveling torsion wave propagating from base to tip.

### Local cross-section rotation during beating measured using gold nanoparticles (GNPs)

To test if a rotation of the local bending plane (torsion) is accompanied by an equal rotation of the axonemal cross-section (twist), we first need to determine both rotations in a common frame of reference, the laboratory frame. There, a rotation of the local bending plane is characterized by Δ*ω*_3D_ (*ϕ, s*), and a local rotation of the axonemal cross-section is characterized by Δ*ω*_GNP_ (*ϕ, s*) (see **Figure 1C**). To measure the Δ*ω*_GNP_(*ϕ, s*), we attached GNPs to the surface of beating axonemes which were then imaged at 5000 fps (**Figure 4**). We determined the projected distance *d*_C_ between the GNP and the axoneme centerline in 2D images with nanometer precision (**Figure 4A**, Materials and Methods). We find that *d*_C_ oscillates at the frequency of the axonemal beat with peak-to-peak amplitudes ranging from 13 to 124 nm depending on the azimuthal positions of the GNPs on the axoneme (see **Figure 4B** for a typical example, and **Figure S9**). Using simulated data, we confirmed that the observed peak-to-peak amplitudes are larger than potential curvature-dependent tracking error with expected magnitude of 8 nm (or less) for curvature values of 0.8 rad/µm (or less) (see **Figure S6E**). To reduce noise, we computed the beat-cycle average *d*_C_(*ϕ*) as function of the beat-cycle phase *ϕ* for each GNP individually, similar to the computation of the average 3D waveform (red line in **Figure 4C**). The change of *d*_C_ during the beat-cycle is indicative of a rotation of the local axonemal cross-section. From the *d*_C_(*ϕ*) profile, we calculate the cross-section rotation angle *ω*_GNP_(*ϕ*) with respect to the laboratory frame, using the known radii of the axoneme (*r*_axoneme_ = 100 nm)^27,28^ and the GNP (*r*_GNP_ = 25 nm) (**Figure 4C**, right axis). This defines the peak-to-peak rotation amplitude Δ*ω*_*GNP*_ of the oscillating rotation angle (**Figure 4D**). Finally, we determine Δ*ω*_GNP_ (s) as function of arc-length *s* by combining results from all GNPs (**Figure 4E**). In the following, Δ*ω*_GNP_ (*s*) and Δ*ω*_3D_(*s*) are used to compare the rotation of the axonemal cross-section to the rotation of the bending plane.

**Figure 4:**
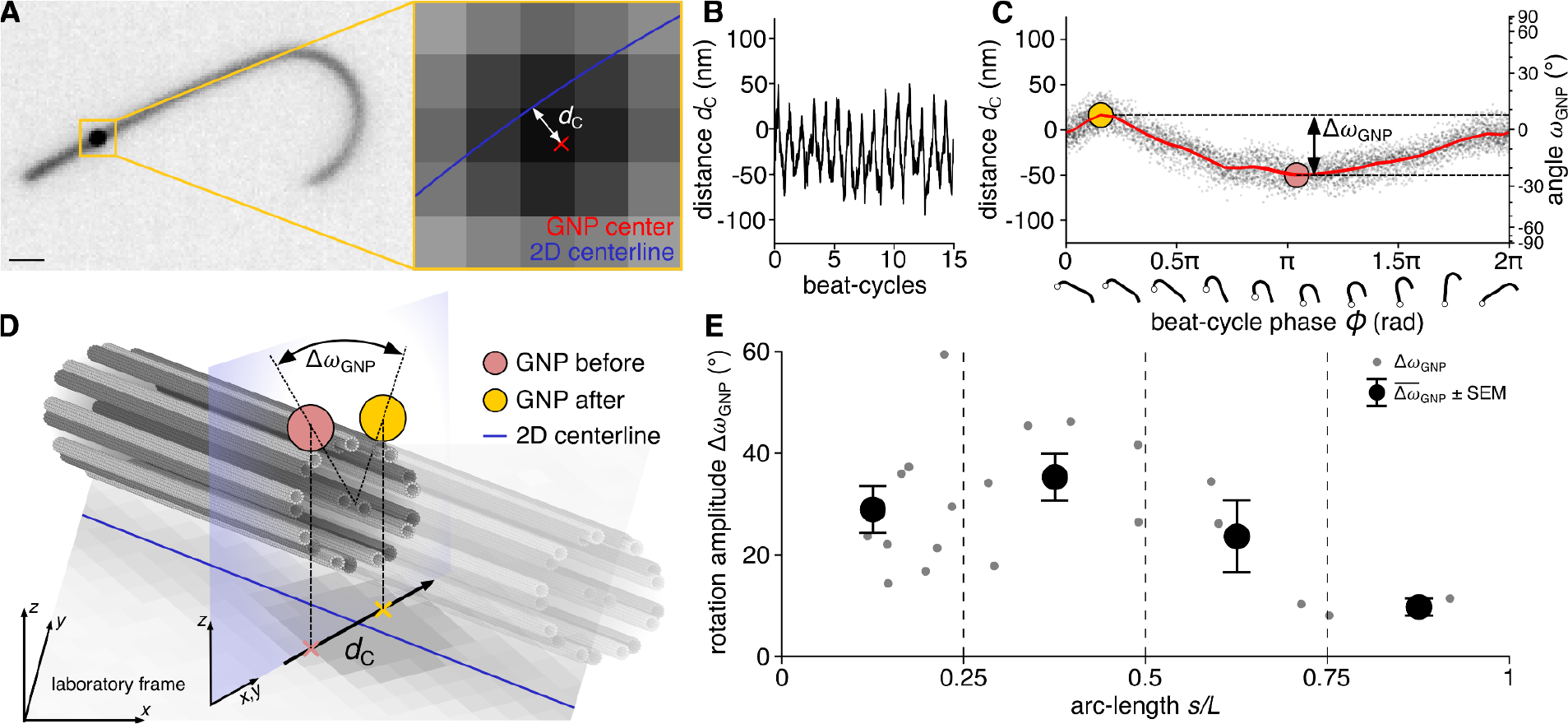
Local cross-section rotation measured using gold nanoparticles (GNPs) attached to beating axonemes. **(A)** High-speed darkfield microscopy image (exposure time 193 µs) of reactivated axoneme with attached GNP (scale bar 1 μm, right panel: zoom-in). We fit a Gaussian model describing the intensity profile of the GNP and the nearby axoneme (Materials and Methods, Table S1) to precisely measure the centerline (blue) and GNP position (red cross) to calculate their projected distance *d*_C_ in the 2D image (pixel size 73 nm). **(B)** Exemplary time-dependent distance to the centerline *d*_C_ (Materials and Methods and **Figure S7**). **(C)** Distance to the centerline *d*_C_ as function of beat-cycle phase *ϕ* (gray dots: pooled data from 67 beat-cycles, red line: phase average), from which the rotation angle *ω*_GNP_ of the axonemal cross-section (in laboratory frame) and its peak-to-peak amplitude Δ*ω*_GNP_ can be computed. **(D)** Visualization of Δ*ω*_GNP_ (*ϕ*). **(E)** Peak-to-peak amplitude Δ*ω*_GNP_ for 19 axonemes, each with one or two GNP attached at different arc-length position (gray filled circles) and averages after binning arc-length position (black filled circles with whiskers, mean ± SEM, bin boundaries indicated by dashed lines).

### Indication for twist-torsion coupling

If twist and torsion are coupled, the measured torsion waves would imply that there are equal twist waves and that the axonemal cross-section and the bending plane would rotate together (**Figure 1C**). We quantify the local bending plane rotation angle *ω*_3D_(*ϕ, s*) from the average 3D waveform and computed the rotation amplitudes Δ*ω*_3D_ (*s*) (red curve in **Figure 5A**, see also **Figure S4A** for map of *ω*_3D_ (*ϕ, s*)). Because we measure Δ*ω*_3D_ (*s*) within the region-of-trust, it provides a lower bound for the rotation amplitude of the bending plane within the beat-cycle. In the scenario of twist-torsion coupling, Δ*ω*_3D_ (*s*) should be equal to Δ*ω*_GNP_(*s*) (blue curve in **Figure 5A**). In the opposite scenario of twist-free torsion, there is no twist and hence, no axonemal cross-section rotation due to twist (**Figure 1D**). However, an attached GNP could still show a rotation with peak-to-peak amplitude Δ*ω*_no twist_ (*s*) relative to the laboratory frame, due to axoneme rolling. We estimated Δ*ω*_no twist_ (*s*) using the measured 3D waveform and a hydrodynamic minimization argument (gray curve in **Figure 5A**, see also **Figure S11** and SI text). When we compare the above-described rotation amplitudes we find that Δ*ω*_GNP_ (*s*) matches Δ*ω*_3D_ (*s*) but not Δ*ω*_no twist_ (*s*) (**Figure 5A**). Specifically, the mean values of Δ*ω*_GNP_ (*s*) and Δ*ω*_3D_ (*s*), calculated for each arc-length bin, are not significantly different (Student’s t-test, 2-sided, unpaired, *α* = 0.05). Likewise, the average difference *d* = Δ*ω*_GNP_ (*s*) − Δ*ω*_3D_ (*s*) within the region-of-trust is not significantly different from zero (**Figure 5B**). In contrast, the rotation amplitude calculated for the twist-free torsion scenario Δ*ω*_no twist_ is consistently lower than the experimental measurement of Δ*ω*_GNP_ (**Figure 5A**). Specifically, the mean values of Δ*ω*_GNP_ (*s*) and Δ*ω*_no twist_ (*s*), calculated for each arc-length bin, are significantly different (Student’s t-test, 2-sided, unpaired, *α* = 0.05). Additionally, we quantify the difference *d* = Δ*ω*_GNP_ − Δ*ω*_no twist_ within the entire beat-cycle which is significantly different from zero (**Figure 5B**). Thus, our data is not consistent with the hypothesis that the 3D waveform is generated without twist but consistent with twist-torsion coupling.

**Figure 5:**
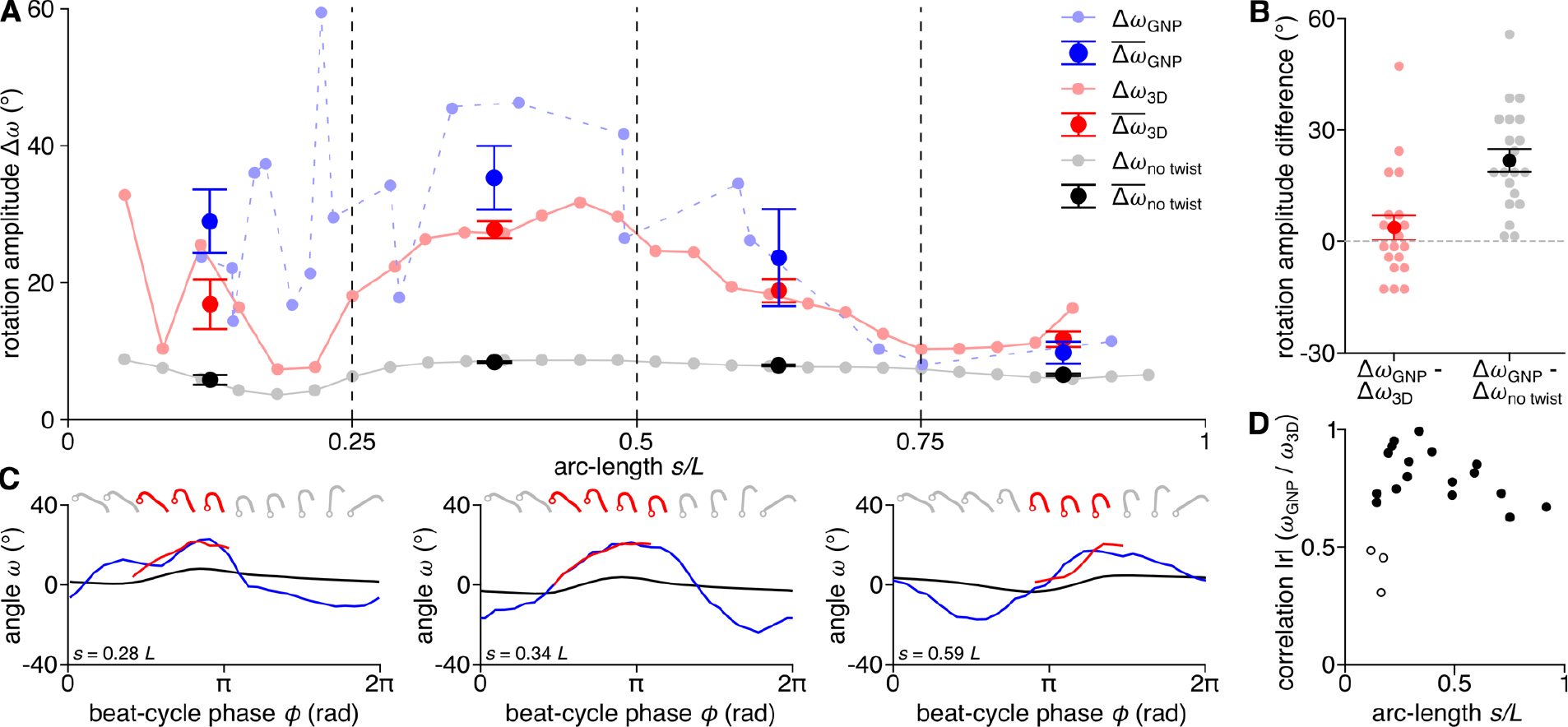
Indication for twist-torsion coupling. **(A)** Rotation amplitudes Δ*ω*_GNP_ (light-blue circles), Δ*ω*_3D_ (rosé circles) and Δ*ω*_no twist_ (gray circles), together with the corresponding means ± SEM in consecutive arc-length bins of length 0.25 *L* (blue, red, black). **(B)** Scatterplot of the differences between Δ*ω*_GNP_ and Δ*ω*_3D_ (measured in the region-of-trust) and between Δ*ω*_GNP_ and Δ*ω*_no twist_ (measured in the entire beat-cycle). These differences are approximately normally distributed (Kolmogorov-Smirnoff test, with Δ*ω*_GNP_ - Δ*ω*_3D_: *p* = 0.41 and Δ*ω*_GNP_ - Δ*ω*_no twist_: *p* = 0.68). The average difference between Δ*ω*_GNP_ and Δ*ω*_3D_ was 3.8 ± 3.4 ° (mean ± SEM, n = 20) and not significantly different from zero (Students t-test, two-sided, *p* = 0.27). The average difference between Δ*ω*_GNP_ and Δ*ω*_no twist_ was 21.8 ± 3.1 ° (± SEM, n = 20), significantly different from zero (Students t-test, two-sided, *p* < 0.001). **(C)** Rotation angles *ω*_GNP_ (*ϕ, s*) (blue line), *ω*_3D_ (*ϕ, s*) (red line, shown for region-of-trust) and *ω*_no twist_ (*ϕ, s*) (black line) for three example arc-length positions (*s* = 0.28 *L*, 0.34 *L* and 0.59 *L*). Since the sign and the offset between the curves are unknown, these were determined by minimizing squared residuals. **(D)** Scatterplot of the un-signed Pearson correlation coefficient |*r*| calculated between *ω*_GNP_ (*ϕ, s*) and *ω*_3D_ (*ϕ, s*) using the MATLAB function *corrcoef*. 17 filled circles indicate significant correlation (*p* < 0.05); three open circles indicate non-significant correlation (*p* > 0.05).

### Temporal correlation of axonemal cross-section rotation and bending plane rotation

If twist and torsion are coupled, not only the peak-to-peak amplitudes of the GNP rotation and the bending plane rotation should agree, but also their dynamics during the beat-cycle. The direct comparison between *ω*_GNP_ (*ϕ, s*) and *ω*_3D_ (*ϕ, s*) within the region-of-trust (**Figure 5C**) shows that their dynamics agree: we computed the un-signed Pearson correlation of *ω*_GNP_ (*ϕ, s*) and *ω*_3D_ (*ϕ, s*) and find a significant correlation for GNP positions above *s* = 0.2 *L* (*p* < 0.05; mean correlation coefficient 0.82, **Figure 5D**). This further supports the hypothesis of twist-torsion coupling. In the basal region (below *s* = 0.2 *L* corresponding to the most proximal 2.5 µm of the axoneme) we also find three GNPs (out of nine GNPs measured at similar location) with non-significant correlations. This may be due to measurement uncertainties in this region where the rotation amplitudes are low. Taken together, the temporal correlation of axonemal cross-section rotation and bending plane rotation further supports the hypothesis of twist-torsion coupling.

## Discussion

### Torsion waves in beating axonemes shape their 3D waveform

We measured the 3D waveform of isolated *Chlamydomona*s axonemes with unprecedented spatio-temporal resolution (32 averaged shapes in a beat-cycle, 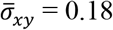 nm and 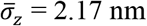) using a combination of high-speed 3D shape reconstruction and beat-cycle averaging. This waveform is made available with this publication (see Extended Source Data). Our approach enables to measure high-resolution torsion profiles and provide rigorous error bounds. Within a beat-cycle, torsion oscillated with an amplitude of 21.9 º/µm (with error below 4.2 º/µm), consistent with an heterochiral torsion wave (combining left and right handedness) propagating from the basal end of the axoneme to its distal tip. Our results are in agreement with earlier studies on reactivated *Chlamydomonas* axonemes ^20^. Torsion generates strongly non-planar shapes. In isolated axonemes, we found that the non-planarity is maximal during the recovery stroke (*ϕ* = *π*) with strong negative torsion in the bend region. This observation is consistent with measurements of the non-planarity in intact *Chlamydomonas* cilia ^29^.

It is known that *Chlamydomonas* cells rotate during swimming, which is important for their phototaxis ^29^. The out-of-plane component of the 3D waveform measured here may contribute to this cell rotation. However, the observation that *Chlamydomonas* cells rotate predominantly during the power stroke of their cilia^30^ (when the beat is mostly planar), and not during the recovery stroke (when we observe strongly non-planar shapes) suggests another relevant contribution, namely the possibility of twist near the axonemal base. More generally, helical navigation due to a non-planar waveforms is also relevant for the chemotaxis of sperm cells ^9^.

### Twist and torsion are coupled in beating axonemes

By comparing the cross-section rotation measured by GNPs attached to the surface of axonemes to the rotation of the local bending plane measured from the 3D shapes, we provide evidence that torsion and twist are coupled. We find that the bending plane and axonemal cross-section rotate together, implying a highly anisotropic bending stiffness of the axoneme which strongly supports the “rigid-bridge hypothesis” - but not the opposite hypothesis of twist-free beating. This result is consistent with the measurements of inter-doublet sliding in reactivated *Chlamydomonas* axonemes using cryo-electron tomography, where sliding amplitudes were low between DMT1-DMT2 as well as between DMT5-DMT6 (coplanar with the bridge) and high normal to the plane of the bridge ^31^. The observed hetero-chiral torsion waves are therefore indicative of equal hetero-chiral twist waves. Our data provide a missing link between torsion (characterizing centerline shapes) and twist (characterizing mechanical deformations).

### Relation to past work

The structural similarities among motile axonemes across species (e.g. the bridge structure) strongly suggest that twist-torsion coupling may be a general feature ^32,33^. To measure the 3D waveforms of beating cilia and flagella, previous studies employed stereographic imaging ^34^, digital in-line holography ^15,35^, multi-focal 2D dark-field microscopy ^16^, defocused 2D bright- and dark-field microscopy ^17,25^, as well as multi-focal phase contrast microscopy ^20^. However, these studies were limited in spatio-temporal accuracy, which made it difficult to reliably quantify torsion. Nonetheless, the consistent observation of non-planar shapes of *Chlamydomonas* cilia ^29^, *Paramecium* cilia ^34^, sperm flagella ^15–17,36^ and Malaria and *Trypanosoma* parasites ^35,37^ indicate the existence of torsion. Twist-torsion coupling now allows to interpret previous torsion measurements as indirect twist measurements. Our observation of twist in beating axonemes is in agreement with twist measurements in fixed axonemes from *Paramecium* ^38^, *Trypanosoma* ^39^ and *Ciona* sperm ^40^ and the observation of the lateral movement of mitochondria during the beat of surface attached quail sperm is consistent with twist waves that we find in *Chlamydomonas* axonemes ^24^.

### Twist generation in the axoneme

Our finding of hetero-chiral twist waves raises the question of how twist is generated. The axoneme has an inherently chiral architecture where dynein motors attached to each DMT exert active forces on the clockwise neighboring DMT when viewed from the base. The activity of the dyneins slides adjacent DMTs. Sliding is restricted at the base. Bending and twist are then generated according to following geometric concepts: (i) If there is a difference in DMT sliding on opposite sides of the axoneme cross-section, the axoneme bends ^41^. (ii) If there is net-sliding, defined as the sum of signed sliding displacements around the circumference of the axoneme, the axoneme twists ^42^. We illustrate these geometric concepts in three examples: (i) If dyneins induce active sliding on only one side of the axoneme, and DMTs on the opposite side are free to slide in the opposite direction, the net-sliding is zero and there is bending but no twist. (ii) If dyneins are active on both sides of the axoneme and induce the same amount of sliding, the net-sliding is non-zero and there is twist but no bending. (iii) If dyneins induce an unequal amount of sliding on opposite sides of the axoneme, there will be both bending and twist. Due to the chiral arrangement of the dyneins that walk towards the microtubule minus ends (i.e. towards the base), the dynein-generated sliding always induces dextral twist.

How sinistral twist is generated is unclear. Potentially, the axoneme might be already sinistrally twisted in the absence of dynein forces. Interestingly, structural sinistral twist was reported for non-motile 9+0 axonemes of human islet cilia ^43^. Sinistral structural components were also reported for motile 9+2 axonemes, such as the central apparatus ^23,44^, or the heads of the radial spokes ^45^. On the other hand, axonemal twist could also be generated by the side-stepping of dyneins ^19^. In gliding assays, it was observed that dyneins can rotate microtubules clockwise when viewed from the minus end ^46,47^. We argue that this rotation would translate into sinistral twist in the axoneme. Taken together, hetero-chiral twist waves could emerge if the axoneme is twisted sinistrally by default and gets periodically unwound and even dextrally over-twisted by dynein-generated sliding.

### Mechanisms of motor control

But how is the activity of dyneins controlled? The precise mechanism of motor coordination driving the axonemal beat remains a matter of debate ^48–56^. Almost all theoretical models assume that motor activity is regulated by mechanical deformations such as curvature, sliding or DMT spacing. A recent theory suggested that twist contributes to regulate dynein activity ^19^. In short, in an axoneme that is both bent and twisted, DMT spacing changes in a different way on opposite sides of the axoneme, which could potentially decrease the distance between dyneins and DMTs and thus regulate their activity. This and other theories prompt precise measurements of time-resolved mechanical deformations of the axoneme such as twist investigated here. Specifically, by providing evidence for twist in bent axonemes, our study presents the first experimental support for a twist-assisted curvature control model as proposed in ref. ^19^. Our quantitative measurements of axonemal twist with oscillations amplitudes of 22 º/µm imply an accumulated twist angle of >16.5 º over distances of 1.5 µm. This is in agreement with the theoretical prediction by ref. ^19^, where an accumulated twist angle of 14 º was sufficient to generate transversal forces strong enough to change the DMT spacing. In conclusion, our data can serve to discriminate between this and alternative models of motor coordination in beating axonemes.

### Summary and Outlook

We provide strong evidence for twist waves in beating *Chlamydomonas* axonemes. This twist generates non-planar 3D waveforms, which can contribute to cell rotation and helical swimming, required for the physiological function of cilia and flagella. Twist is a dynamic mechanical deformation generated by active motor forces. This deformation could regulate dynein motors during axonemal beating. Together, this would put twist at the heart of a proposed mechano-dynamical feedback loop, where motor activity deforms the axoneme, while the resulting mechanical deformation may regulate motor activity.

## Materials and Methods

### Experiments

#### Isolation and reactivation of Chlamydomonas axonemes

Axonemes from *Chlamydomonas reinhardtii* wildtype cells (cc-125 mt+) were isolated and reactivated following the experimental procedures described in ^57^. All reagents were purchased from Sigma Aldrich, unless stated otherwise. In brief, cells were grown in TAP medium under constant illumination by LED-light pads (light therapy lamp HST-001, 10000 LUX, 10W, 4500K) and air-bubbling at 22-24°C for 3-4 days to a final density of 3-7 ⋅ 10^6^ cells mL^−1^. Cilia were isolated by the dibucaine procedure, separated from the cell bodies by centrifugation (2400g, 25% sucrose cushion) and demembranated in HMDEK (30 mM HEPES-KOH, 5 mM MgSO_4_, 1 mM DTT, 1 mM EGTA, 50 mM potassium acetate, pH 7.4) augmented with 1% (v/v) IGEPAL and 0.2 mM Pefabloc SC. The membrane-free axonemes were resuspended in HMDEKP (HMDEK with 1% (w/v) polyethylene glycol, 20 kDa) with 30% sucrose, 0.2 mM Pefabloc added and stored at -80°C. Prior to reactivation, axonemes were thawed at room temperature, then kept on ice for at most 3 hours. The reactivation was performed in flow chambers (with a depth of 100 µm) built from easy-cleaned glass and parafilm ^57^. For the experiments with gold nanoparticles (GNPs), 1 µL of thawed axonemes was mixed with 1 µL of gold nanoparticles (GNP) solution (diameter 50 nm, 3.5 ⋅10^10^ ml^-1^) and incubated for 10 min on ice. Axonemes with or without GNPs were diluted in HMDEKP reactivation buffer containing 1 mM ATP and an ATP-regeneration system (1 unit/ml creatine kinase, 5 mM creatine phosphate) used to maintain the ATP concentration. The axoneme dilution was infused into a glass chamber, which was blocked with casein solution (from bovine milk, 2 mg/mL) for 10 min. Prior to imaging, the flow chamber was sealed with twinsil-speed (picodent) to avoid evaporation. The sample was equilibrated on the microscope for 5 min before data was collected for a maximum time of 60 min.

### Imaging of axonemes

Reactivated axonemes were imaged by darkfield microscopy set up on an inverted Nikon ECLIPSE Ti2-E microscope, using a Nikon 100x iris oil (0.5 – 1.2 NA) lens in combination with a 1.5x or 1.0x tube lens and a Nikon 1.3 NA oil condenser. Images were recorded with a pco.dimax CS4 high-speed camera operated by NIS-Elements Imaging Software (Nikon). *Z*-scan images for 3D calibration were recorded with a pco.edge 4.2 camera. In both cases, the illumination was performed by a Sola SE2 light engine (Lumencor) combined with a 496 LP filter (Semrock, Brightline). Movies of reactivated axonemes with GNP were recorded with the 1.5x tube lens and an NA setting of 1.1 - 1.2 on the objective at a frame rate of 5000 fps. In total, we recorded more than 150’000 image frames from a total 20 GNPs (5000-10’000 images each) attached to 19 beating axonemes (one with two GNPs), corresponding to a total of more than 1750 beat-cycles. For the reconstruction of the 3D waveform, we recorded movies at 1000 fps using a 1.0x tube lens and an objective NA of 1.0. For 3D reconstruction, we first focused below the axoneme, so that the axoneme never entered the focal plane. We then recorded a total of 53’000 defocused images from a total 17 axonemes (3000-5000 each), corresponding to a total of more than 3750 beat-cycles.

### Data analysis

#### Calibration of defocused darkfield microscopy and 3D shape reconstruction

We reconstructed the 3D shape of the axonemal centerline in single defocused images of beating axonemes from the *xy*-position measured by 2D filament tracking and the *z*-position derived from the full-width-at-half-maximum (FWHM) of intensity distributions measured normal to the long axis of the axoneme. All analysis was done with MATLAB (Version R2020a) unless stated otherwise. Specifically, we subtracted the static background (the median intensity in each pixel calculated for the entire movie) using Fiji. We then determined the 2D centerline using the filament tracking software FIESTA ^26^. We interpolated the 2D centerline using a smoothing spline fit (MATLAB) to gain equidistant points with a spacing of 110 nm along the centerline. Subsequently, we determined the intensity distributions along line-scans normal to local tangent of the 2D centerline at each control point of the 2D axoneme centerline and measured the local full-width-at-half-maximum (FWHM) by Gaussian fitting.

To relate the FWHM to an absolute z-position with respect to the focal plane, we performed the following calibration. We non-specifically immobilized axonemes (without ATP) on a di-chloro-dimethyl-silane (DDS) coated glass coverslip. Subsequently, we recorded a z-scan with a range of ±10 µm relative to the z position at which the axoneme was in focus. To gain a calibration curve we measured the FWHM as a function of the distance to the focal plane (z-position where FWHM is minimal, calibration curve shown in **Figure S2E**). To account for the refractive index mismatch between the immersion oil and water (reactivation buffer), we corrected our calibrated z-positions by a factor of 1.137, which is the ratio of the both refractive indexes (n_oil_ = 1.515, n_water_ = 1.333). We estimated the localization uncertainty (standard error) of 3D shapes obtained in single images as *σ*_*xy*_ = 7.3 nm and *σ*_*z*_ = 33.7 nm (SI and **Figure S1**)

#### Beat-cycle phase and beat-cycle averaging

To determine the beat-cycle phase *ϕ* of a given shape, we fitted the arc-length profile of the 2D tangent-angle (time average subtracted) with a sinusoidal function (see sup Supplementary Information and **Figure S2A-D**). By this definition, the recovery-stroke included shapes with the beat-cycle phases [0.5π ≤ *ϕ* < 1.5π]. The remaining shapes belonged to the power-stroke (**Figure S2D**). For details on the determination of the beat-cycle phase, see Supplementary Information and **Figure S2A-D**. To calculate a high-resolution 3D waveform of reactivated *Chlamydomonas* axonemes, we used beat-cycle averaging. To do this we subdivide the beat-cycle into 32 bins (with equal width of Δ*ϕ* = π/16, and group shapes of similar beat-cycle phase. For each bin, we calculated the average shape with 30 points along the arc-length (Δ*s*_2D_ = *L*/30). To do this we: (i) averaged the profiles of the 2D curvature along arc-length *к* _2D_ (*s*_2D_) and the profiles of the z-position along arc-length *z*(s_2D_) of all recorded axoneme shapes, and (ii) calculated the average 2D position *x*(*s*_2D_) and y(*s*_2D_) from the average curvature *к*_2D_(*s*_2D_) by integration. We obtained the averaged *x, y, z* positions along the 2D centerline in time (for each beat-cycle bin), which together comprise the average 3D waveform. For details on the averaging method, see Supplementary Information and **Figure S2E-J**. We estimate the localization uncertainty of the average 3D shapes obtained through beat-cycle averaging as *σ*_*xy*_ = 0.18 nm and *σ*_*z*_ = 2.17 nm (SI and **Figure S2K**).

#### Measurement of the distance between gold nanoparticles and the axoneme centerline

We precisely measured the distance between gold nanoparticles (GNP) and axoneme 2D centerline *d*_C_ in each recorded image. To do this, we used FIESTA ^26^, to roughly measure the position of the axoneme-bound GNP in the image. Using this position, we selected a region of interest of 25x25 pixels, which contained the GNP in the center. Within this box, we fitted a 2D intensity model, which was the sum of a symmetric 2D gaussian function and a gaussian wall function. The center of the symmetric 2D gaussian described the center position of the GNP while the centerline of the gaussian wall described the axoneme centerline. We used both to calculate the normal distance between the GNP and the axoneme centerline, which we called distance to centerline *d*_C_. We subtracted an orientation-dependent systematic error from the measured *d*_C_ (detailed in the Supplementary Information and **Figure S6**). To calculate the average *d*_C_ as a function of the beat-cycle phase for a single GNP, we performed beat-cycle averaging as described above. We used the average *d*_C_(*ϕ*) to calculate the angular position of the GNP in the cross-section as observed from the laboratory coordinate system *ω*_GNP_. Note that since *d*_C_ is only a two-dimensional projection, we cannot measure whether the GNP is attached above or below the axoneme, which would change the sign of *ω*_GNP_. We quantified the local cross-section rotation by the peak-to-peak amplitude Δ*ω*_GNP_. We determined the approximate point of GNP attachment along the arc-length of the axoneme via a segmented line measurement tool (Fiji) and normalized this distance by the axoneme length.

#### Calculation of the 3D curvature, torsion and ω_3D_

A space curve is parameterized by arc-length s with three unit-vectors of the Frenet-Serret Frame: tangent vector T, the principal normal vector N and the binormal vector B. These three vectors are mutually orthogonal B = T × N. If the curvature is non-zero, B is the normal vector of the osculating plane that is spanned by T and N. Here the curvature *к* of the space curve can be calculated as the quotient of arc-length derivative of T and N. The torsion *τ* of the space curve can be calculated as the quotient of arc-length derivative of B and N.

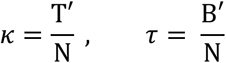

The measured 3D shapes represent discretized space curves for which we calculate approximations for the unit vectors of the Frenet-Serret frame 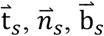. We approximate (i) the local tangent vector at a given point with the mean of the two adjacent secant vectors of the discretized space curve, (ii) the local binormal vector as the cross-product of the two adjacent tangent vectors and (iii) the local normal vector as the cross-product of the binormal vector and the additive inverse of the tangent vector. We require that the *z*-component of the binormal vector is always positive. Thereby, we can define a unique sign of the curvature for the entire 3D shape. Using the approximation of the Frenet-Serret frame, we calculate torsion and curvature as the arc-length derivative of the signed rotation angle of the local bending plane (osculating plane) *ω*_*τ*_ and the signed in-plane rotation angle Ψ respectively. Additionally, we obtain the local bending plane orientation with respect to the laboratory coordinate system *ω*_3D_ as the angle between the normal vector and the *xy*-plane. We present these 3D shape parameters as a function of arc-length and beat-cycle phase *κ* (*ϕ, s*), *τ*(*ϕ, s*) and *ω*_3D_ (*ϕ, s*) (**Figure 3D, E** and **Figure S4A**).

#### Sign conventions

We defined the sign of the curvature of the static waveform shape component ^58^ as positive (as seen from the axoneme base). Segments of curves that have a dextral handedness show positive torsion and segments with sinistral handedness show negative torsion. The sign of *ω*_GNP_ and *ω*_no twist_ was chosen to oppose the sign of the *z*-component of the normal vector, from which it was calculated (**Figure S4A**). The sign of *d*_C_ is defined with respect to the axoneme orientation (SI). The sign of the measured *ω*_GNP_ is arbitrary for individual GNPs since we do not know on which side of the axoneme the considered particle was bound. To compare the temporal dynamics of *ω*_GNP_ (*ϕ, s*) and *ω*_3D_ (*ϕ, s*) in **Figure 5D** we therefore use the *un-signed* Pearson correlation coefficient (calculated using the MATLAB function *corrcoef*) as sign of the correlation depends on the whether the GNP was bound to the upper or lower side of the axoneme, which is unknown. A method that could be used to define a global sign of *ω*_GNP_ is suggested in the Supplementary Information (**Figure S9**).

#### Torsion: measurement uncertainty and the choice of the region-of-trust

To estimate the uncertainty of the torsion measurement, we used a bootstrapping approach. Our experimental dataset contained 53’000 shapes (from 17 axonemes). From those shapes, we drew 1’000 sets of 53’000 shapes with replacement. For each set, we calculated the average 3D shape as well as the torsion *τ*(*ϕ, s*). The standard deviation over all 1’000 computed torsion maps was used as an estimator of the measurement uncertainty of the torsion. To determine the region-of-trust of the torsion measurement, we used the argument that the normal (and binormal) vector of the Frenet-Serret frame is ill-defined in regions of low curvature. Since torsion is calculated using those vectors, the variability of torsion increases with decreasing curvature (**Figure S3**). Based on this dependence, we defined a conservative region-of-trust in which the variability is approximately constant, which is the case for values of local curvature of |*κ* | > 0.4 rad/µ*m* and above. The same region-of-trust was used for the rotation angle *ω*_3D_ of the local bending plane relative to the laboratory frame.

## Acknowledgements

We thank Joe Howard for critically reading the manuscript and all members of the Diez lab for fruitful discussions. Funding for this work was provided by the Dresden International Graduate School for Biomedicine and Bioengineering (DIGS-BB), Technische Universität Dresden (TUD) and the Max Planck Institute of Molecular Cell Biology and Genetics Dresden (MPI-CBG).

## Data availability statement

All imaging data and positional information extracted from the images are available on the data repository Zenodo (https://doi.org/10.5281/zenodo.10513932).

## Code availability statement

All code and evaluation scripts are available from the corresponding authors on request.

