## Supplementary Information for "Twist - torsion coupling in beating axonemes"

**3D defocused darkfield microscopy**

We used a standard darkfield-microscopy setup based on a Nikon Ti2 microscope. As light-source, we use the Sola2 light engine (lumencore) together with a 496 nm filter. We illuminate the sample using a 1.4 NA Nikon oil condenser and capture the scattered light with a 100x Nikon iris oil objective with variable NA (0.5 – 1.2), set to NA 1.0. We record images with a PCO.dimax CS4-M9-GE high-speed camera with an exposure time of 1 ms (Figure 2B).

Defocused darkfield microscopy images of isolated and reactivated axonemes were recorded by defocusing below the sample (focal plane gray dotted line in, Figure 2C). The reactivated axonemes we imaged were always close to the lower surface of the flow chamber. We tracked the 2D shapes in these images using the filament tracking software package FIESTA (Ruhnow, Zwicker, and Diez 2011) yielding *x-* and *y-*coordinates along the 2D arc-length $s_{2D}$, where $s_{2D}$ = 0 and $s_{2D}$ = *L* correspond to the proximal and the distal end of the axoneme, respectively (where the proximal points in the direction of swimming). To obtain the axial (*z*) position of the axoneme, we measured the axoneme intensity distribution through line-scans normal to the 2D projection of the tracked axoneme centerline at each arc-length position. Two exemplary line-scans show positions where (1) the axoneme was highly defocused (red line, Figure 2B) and (2) where the axoneme was close to the focal plane (blue line, Figure 2B). We fitted the respective intensity profiles with Gaussian functions to determine the full-width-at-half-maximum (FWHM), which was low where the axoneme was close to the focal plane and high where it was further away from the focal plane (Figure 2E). Using a *z*-scan of immobilized axonemes, we calibrated the observed FWHM at given distance to the focal plane. We obtained a calibration curve by performing a cubic spline fit with the MATLAB function ‘fit‘ (fit-type ‘smoothingspline’ with smoothing parameter 0.9, green line, Figure 2E). Using this calibration curve, we determine the local *z*-positions of shapes of beating axonemes from the measured FWHM obtained from line-scans.

**Error estimation for 3D shapes from defocused darkfield microscopy**

To estimate the axial localization uncertainty, we measured the residuals between *z*-position $z_{\mathrm{raw}}\left( s_{2D} \right)$ as function of 2D-arc-length $s_{2D}$ and fitted curves, similar to (Mojiri et al. 2021). Here, we fitted $z_{\mathrm{raw}}\left( s_{2D} \right)$ of all recorded images with the MATLAB function ‘fit‘ using the fit-type ‘smoothingspline’ (Figure S1A, $s_{2D}$ was calculated from the xy-position in the image). Using this option for fitting, a smoothing parameter (sm) was required to set the sensitivity of the resulting fit function with respect to the variance in the data. To find this optimal smoothing parameter, we calculated the standard deviation $\sigma_{z}$ of the distribution of *z*-residuals as function of the smoothing parameter and determined the position where the absolute slope of this curve was minimal (Figure S1C). We found the optimal smoothing parameter to be sm = 0.72, and used this to smoothen $z_{\mathrm{raw}}\left( s_{2D} \right)$ and obtain $z_{sm=0.72}\left( s_{2D} \right)$.$Then the$ standard deviation of the residual distribution was $\sigma_{z}$ = 33.8 nm (Figure S1B). We found that 95% of the data-points constituting all measured axonemal shapes were in the range of 0.70 µm - 1.35 µm above the focal plane (Figure S1D, E). This shows that the *z*-component of the 3D waveform varies by about 0.65 µm for axonemes swimming near the surface of a coverslip. Additionally, the lower boundary of this range indicates that during recording, the focal plane was located about 0.7 µm below the surface towards which the axonemes were swimming. Thus, within the admissible range of our calibration curve (0.7 µm - 1.35 µm above the focal plane) we find a *z*-dependent error $\sigma_{z}$ in the range of 29 nm - 45 nm (Figure S1E).

To estimate the technical lateral position error $\sigma_{xy}$, we rely on the fit-error estimation of FIESTA. The estimated fit-errors in 2D for all axonemes are approximately normally distributed (fit of histogram with normal distribution: R^2^ = 0.98) with an average fit error of 7.3 nm and a standard deviation of 2.2 nm.

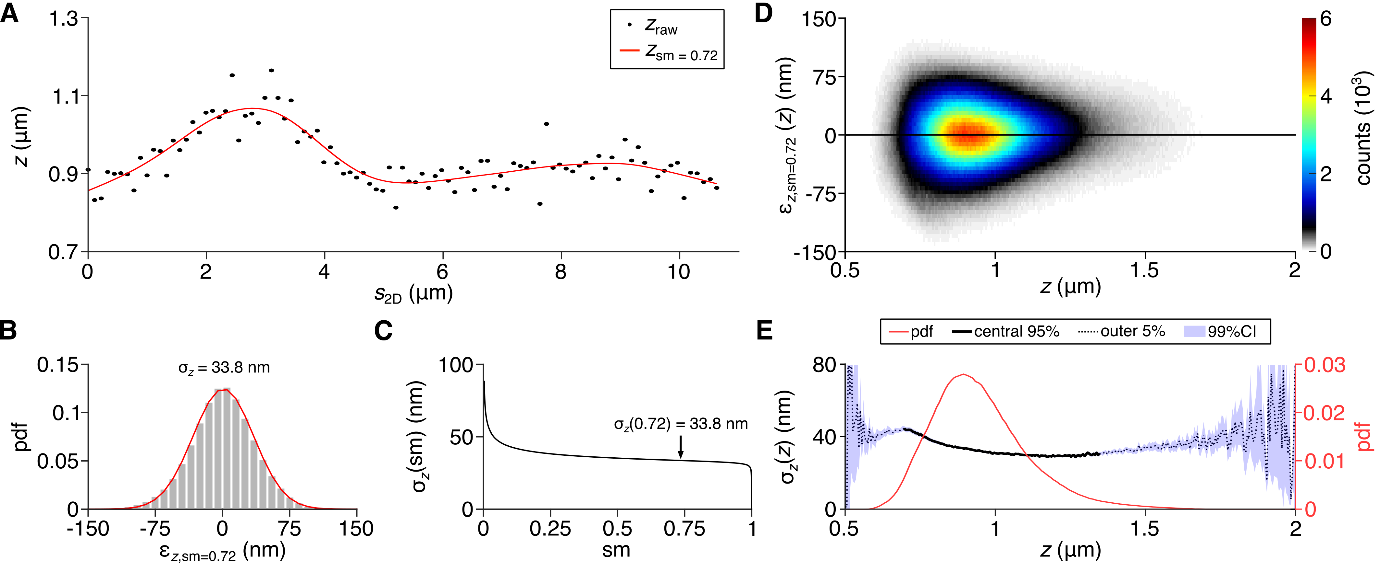

**Figure S1: *Z*-error estimation for 3D shapes recorded with 3D defocused darkfield microscopy.** **(A)** Measured *z*-position (gray points) along the 2D arc-length $z_{\mathrm{raw}}\left( s_{2D} \right)$ of a single recorded image with fitted smoothing spline curve $z_{\mathrm{sm}}\left( s_{2D} \right)$ for smoothing parameter sm = 0.72 (red line). **(B)** Histogram of the residuals $\epsilon_{z, sm = 0.72}$ between $z_{\mathrm{raw}}\left( s_{2D} \right)$ and $z_{\mathrm{sm}}\left( s_{2D} \right)$ for all recorded ≈$6\cdot{10}^{6}$ data-points (n = 17 axonemes in 53’000 images, normalization: probability density function (pdf)). By fitting a Gaussian function (red curve), we obtain $\sigma_{z}$ = 33.8 nm for sm = 0.72. **(C)** $\sigma_{z}$ as function of smoothing parameter sm; the optimal sm = 0.72 is located where this curve’s absolute first derivative is smallest. **(D)** Heat map of residuals $\epsilon_{z}$ as function of the respective *z*-position on the smoothing spline curve for all $\approx6\cdot{10}^{6}$ data-points (sm = 0.72, bin-widths: 5 nm for $\epsilon_{z}$, 5 nm for *z*). **(E)** Probability density function of residuals $\epsilon_{z}$ (pdf, red line, bin-width = 0.01 μm). Relationship between measurement error $\sigma_{z}$ and *z*-position ($sm=0.72$) is shown for the central 95% of the pdf (solid black curve) as well as for the outer 5% (dotted black curve). The 99% confidence interval (CI) of $\sigma_{z}$(*z*) was calculated from the output of the fit function.

**Beat-cycle phase**

The periodic shape changes of beating axonemes can be described as a bending wave traveling from the proximal to the distal end. This periodic shape change can be parameterized by a 2π-periodic phase or clock variable $\phi$. To determine $\phi$, we measure the tangent angles along the 2D arc-length $\psi_{2D}(s_{2D},t)$ and subtract the tangent angle at the basal end, obtain $\psi_{2D}^{'}(s_{2D},t)$ (Figure S2A, B). For each axoneme, we calculate the time-averaged tangent angle as function of 2D arc-length $\left\langle\psi_{2D}^{'}(s_{2D},t) \right\rangle_{t}$ (Figure S2B, red curve). The subtraction of this time-average yields the dynamic component of the waveform (Geyer et al. 2016) (Figure S2C, black curves). We fit these resulting profiles with a sinusoidal function $A \sin\left( \frac{2\pi s_{2D}}{L}-\phi+\pi\right)$ (see Figure S2C). This allows us to quantify the progression of a traveling wave profile (with wavelength approximately equal to the length *L* of the axoneme) as function of the beat-cycle phase $\phi$. Using this method, a $\phi$-value can be assigned to every recorded shape, which replaces the extrinsic time dimension with this intrinsic shape-derived clock variable $\phi$.

Traditionally, the beat-cycle of *Chlamydomonas* cilia is divided into a so-called power stroke, during which both cilia move towards the cell body, propelling the cell forward, and a recovery stroke, during the cilia move backward to a position in the front of the cell. In our parameterization, the shapes of the power-stroke correspond to phase-values between [$\phi$ < 0.5π; 1.5π ≤ $\phi$] (Figure S2D). The shapes of the recovery-stroke correspond to phase-values between [0.5π ≤ $\phi$ < 1.5π] (Figure S2D).

**Beat-cycle averaging**

We use beat-cycle averaging to calculate average 3D shapes for each phase of the beat-cycle, which provides a high-precision measurement of the 3D waveform of the beating *Chlamydomonas* axoneme. This averaging method relies on averaging 3D shape information of many images in which axonemes are in the same phase of the beat-cycle. For this, we divide the beat-cycle into 32 bins of beat-cycle phase with equal width of $\Delta\phi$ = π/16 (Figure S2E) and sort recorded shapes into these bins. For example, the axoneme shape in Figure S2A corresponds to a phase of $\phi$ = 0.05π. Therefore, this shape falls into the first bin of the beat-cycle [0 ≤ $\phi$ < π/16] (Figure S2E). We calculate an average 3D shape for each of these bins. The raw 3D shape information is obtained from single images of defocused darkfield microscopy and comprises the axial position $z(\phi,s_{2D})$ and 2D curvature $\kappa_{2D}(\phi,s_{2D})$. Here, we rely on curvature as a measure for the 2D shape because it is independent of position and orientation of axonemes in the *xy*-plane. The values of $z(\phi,s_{2D})$ and $\kappa_{2D}(\phi,s_{2D})$ are measured at each point along the 2D shape with a spacing of ${\Delta s}_{2D}$ = 110 nm (pixel size). Between images and between axonemes, the number of points along the arc-length varies (average ≈ 113 points in 53’000 images). To superimpose $z(\phi,s_{2D})$ and $\kappa_{2D}\left( \phi,s_{2D} \right)$ of multiple images with the different number of points, we use a down-sampling approach based on the weighted arithmetic mean to reduce the number of points along the normalized 2D arc-length to exactly n = 30 (Figure S2F). Here, the weights for each value in the raw data are determined by the overlap of the corresponding arc-length interval with the new 30-point 2D arc-length binning.

As all axonemes were recorded independently, the average axial positions (average distance to the focal plane) may differ between axonemes. Thus, the axial positions $z(\phi,s_{2D})$ of each axoneme needed to be corrected by subtracting the axoneme-specific total average *z*-position, which was calculated as the average *z*-position along the arc-length and throughout all recorded images of an axoneme.

The 3D shape information of every single recorded image was binned into a grid with 30 bins along the arc-length and 32 bins throughout the beat-cycle (Figure S2G). To determine the average shape, we calculate the average of the *z*-position and of $\kappa_{2D}$in every bin. All bins along the 2D arc-length for the beat-cycle phase bin [0 ≤ $\phi$ < π/16] are displayed in Figure S2H and Figure S2I. After calculating the average in each individual bin, we obtain the average *z*-position as function of 2D arc-length $\left\langle z(s_{2D}) \right\rangle$ (red line in Figure S2H) as well as the average $\kappa_{2D}$as function of 2D arc-length $\left\langle\kappa_{2D}(s_{2D}) \right\rangle$. By assuming a point spacing of ${\Delta s}_{2D}$ = 414 nm along the 2D arc-length, which we derive from the average axoneme length (≈ 12.4 μm) and the number of data-points along the arc-length (n = 30), we reconstruct the average 2D shape of the axoneme in [0 ≤ $\phi$ < π/16] from $\left\langle\kappa_{2D}(s_{2D}) \right\rangle$ (orange shape in *z* = 0 plane in Figure S2J). The average *z*-position as function of the 2D arc-length $\left\langle z(s_{2D}) \right\rangle$ provides the axial position information for the average shape in [0 ≤ $\phi$ < π/16] (red lines in Figure S2J). The result is a 3D centerline, which represents the average 3D shape of the beating axoneme in the interval [0 ≤ $\phi$ < π/16] (Figure S2G, blue line). As the *z*-component is comparably small, the distances between the 3D points $\Delta s$ remain approximately constant with $\Delta s$ = 415.5 ± 2.0 nm (distances between all points of average 3D shapes of all beat-cycle bins, mean ± standard deviation, n = 928). Using this method for all bins, we obtain the average 3D shapes throughout the beat-cycle.

**Error estimation for the shapes of the average 3D waveform**

Assuming that the data-points in each bin of the 30x32 grid (Figure S2G) are approximately normally distributed, the standard error of the mean (SEM) can be used to estimate a confidence interval. The confidence interval determines the range in which the true mean (true position of the axoneme centerline) is located with a given confidence level. This range scales with the SEM, which provides a measure for the positional uncertainty of individual data-points of the 3D shape. We observe that the *z*-uncertainty $\sigma_{z}(\phi,s_{2D})$ranges from 1.29 nm to 5.93 nm, with an average of $\left\langle\sigma_{z} \right\rangle_{s,\phi}$ = 2.17 nm (Figure 2SK, lower panel). We find that $\sigma_{z}$is correlated with the respective average *z*-position (r = 0.82). The *z*-uncertainty $\sigma_{z}$ arises from variability of *z*-positions in a given bin of the 30x32 grid. This variability is due to measurement noise, variability between beat-cycles and variability between axonemes. Because the measurement noise shows a negative correlation with the respective average *z*-position, we can conclude that the $\sigma_{z}$ is dominated by variability between beat-cycles and between axonemes, i.e. the inherent variability of the 3D waveform.

To estimate the lateral positional uncertainty of the 3D shape $\sigma_{xy}(\phi,s_{2D})$, we apply an approach that was previously reported in (Gittes et al. 1993). There, the authors used the geometrical relation between the local positional error (2D) and the error of the local tangent angle, which in turn is related to the local error of the curvature. To apply this concept to our case, we calculate the error of curvature $\sigma_{\kappa_{2D}}(\phi,s_{2D})$ in each bin of the 30x32 grid (Figure S2G). Similar to the estimation of the axial positional uncertainty, we use the SEM of $\kappa_{2D}(\phi,s_{2D})$ in each bin to characterize the local uncertainty of the mean curvature that serves as an estimate for the error. By multiplying $\sigma_{\kappa_{2D}}(\phi,s_{2D})$ with the *xy*-point-to-point spacing of the average shape ${\Delta s}_{2D}$ = 415.5 nm, we calculate the error of the 2D tangent angle from the experimental data. This allows us to estimate the lateral positional uncertainty in 2D $\sigma_{xy}$ with the following formula

$$\sigma_{xy}(\phi,s_{2D})\approx\tan\left( \sigma_{\kappa_{2D}}\left( \phi,s_{2D} \right) {\Delta s}_{2D} \right) \frac{{\Delta s}_{2D}}{2}$$

We find that $\sigma_{xy}$ ranges from 0.05 nm to 0.38 nm, with an average of $\left\langle\sigma_{xy} \right\rangle_{s,\phi}$= 0.18 nm (Figure 2SK, upper panel). We observe a weak correlation of $\sigma_{xy}$ with the average curvature in each bin of the grid (r = 0.61).

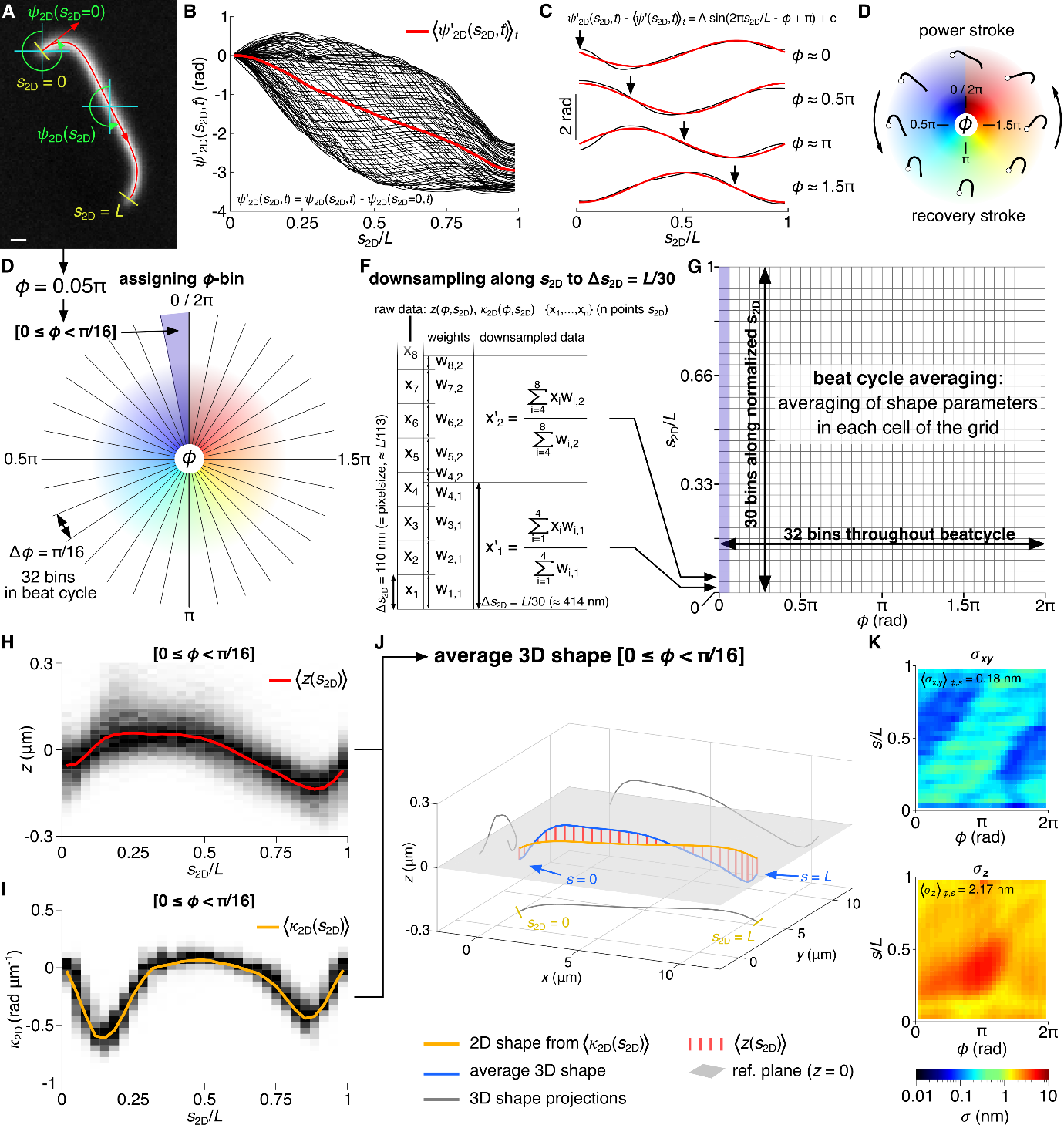

**Figure S2: Beat-cycle phase determination and beat-cycle averaging. (A)** Defocused high-speed darkfield microscopy image of a reactivated axoneme with 2D centerline (red line). Displayed are two 2D tangent vectors (red arrows) and respective 2D tangent angles $\psi_{2D}$ at the basal end ($s_{2D}$= 0) and at approximately $s_{2D}$ = *L*/2. $\psi_{2D}$ is measured with respect to the field of view. Scale bar = 1 μm. **(B)** Exemplary 2D tangent angle as function of 2D arc-length profiles of one beat-cycle (n = 90 curves, from movie recorded at 5’000 fps) where the basal tangent angle has been subtracted (${\psi'}_{2D})$. The time-averaged tangent angle as function of 2D arc-length $\left\langle\psi_{2D}^{'}(s_{2D},t) \right\rangle_{t}$ of a whole time-series is highlighted (red line, whole time series: 5’000 images, 65 beat-cycles). **(C)** The time-average-subtracted 2D tangent angle as function of arc-length (black curves) characterizes the traveling bending wave of the beating axoneme. To calculate the phase of the beat-cycle $\phi$, these curves are fitted with a sinusoidal function $A \sin\left( \frac{2\pi s_{2D}}{L}-\phi+\pi\right)$ (red curve). For this fit, the fit parameters were restricted to the intervals [0 ≤ *A* < ∞] and [0 ≤ $\phi$ < 2π]. **(D)** The recovery-stroke of the beat-cycle includes all shapes in the range [0.5π ≤ $\phi$ < 1.5π], while the power-stroke comprises the remaining shapes. **(E)** For beat-cycle averaging, the recorded images are binned using the beat-cycle phase $\phi$. The beat-cycle is subdivided into 32 bins with equal width of $\Delta\phi$ = π/16. The shape shown in panel A corresponds to a phase angle of $\phi$ = 0.05π and thus is sorted in the first of the 32 bins [0 ≤ $\phi$ < π/16]. **(F)** The raw values of $z(\phi,s_{2D})$ and $\kappa_{2D}(\phi,s_{2D})$ are measured at each point along the 2D shape with a spacing of ${\Delta s}_{2D}$=110 nm (pixelsize). Between images and between axonemes, the number of points along the arc-length varies. For beat-cycle averaging, these arc-length profiles are down-sampled to n = 30 points along the normalized 2D arc-length by using a weighted arithmetic mean (weights are equal to the overlap between arc-length bins corresponding to raw data with spacing ${\Delta s}_{2D}$ and down-sampled data with spacing ${\Delta s}_{2D}$) **(G)** The $\phi$-binned and down-sampled measurables for all recorded images are superimposed in a grid that spans 30 bins along the arc-length and 32 bins throughout the beat-cycle (average *z*-position along $s_{2D}$ and throughout all images for each axoneme is subtracted for each axoneme). **(H)** 2D histogram of combined $z(\phi,s_{2D})$ measurements of all images of all axonemes, where [0 ≤ $\phi$ < π/16] (first bin in beat-cycle, N = 54’330 data-points in total, 1’811 data-points per arc-length bin of width ${\Delta s}_{2D}$). The average *z*-value in each bin along $s_{2D}$ is calculated and yields the average $\left\langle z(s_{2D}) \right\rangle$ (red line). **(I)** 2D histogram of combined $\kappa_{2D}(\phi,s_{2D})$ measurements of all images of all axonemes, where [0 ≤ $\phi$ < π/16] (first bin in beat-cycle, N = 54’330 data-points total, 1’811 data-points per arc-length bin of width ${\Delta s}_{2D}$). The average $\kappa_{2D}$-value in each bin along $s_{2D}$ is calculated and yields the average $\left\langle\kappa_{2D}(s_{2D}) \right\rangle$ for $\phi$-values in the given range (orange line). **(J)** The average 3D shape (blue line) is assembled from the reconstructed 2D shape (orange line, calculated from $\left\langle\kappa_{2D}(s_{2D}) \right\rangle$ with ${\Delta s}_{2D}$ = 414 nm) and the average *z*-position $\left\langle z(s_{2D}) \right\rangle$ (red lines)). The reference plane (*z* = 0) and the projections of the average 3D shape to each coordinate plane of the laboratory frame (gray lines) are displayed. **(K)** Maps of estimated localization errors $\sigma_{xy}$ and $\sigma_{z}$ of the average 3D shapes, calculated from the standard error of mean in each bin of the grid (G). $\sigma_{xy}$ is derived from the standard error of the mean of the 2D curvature in each bin of the grid by using the error estimation according to (Gittes et al. 1993). The average localization errors amount $\left\langle\sigma_{xy} \right\rangle_{s,\phi}$ = 0.18 nm and $\left\langle\sigma_{z} \right\rangle_{s,\phi}$ = 2.17 nm.

**Calculation of torsion error and threshold curvature for the region-of-trust**

We use the Frenet-Serret frame of the shapes of the average 3D waveform to calculate torsion $\tau(\phi,s)$ and 3D curvature $\kappa(\phi,s)$, which results in maps of these shape parameters, which are a function of arc-length and the beat-cycle (Figure 3D, E). Torsion quantifies the incremental rotation of the bending plane along the arc-length of the axoneme. To measure this bending plane rotation along the arc-length, we calculate the orientation of the local bending plane in the axoneme reference frame. Additionally, we calculate the orientation of the bending plane with respect to the laboratory coordinate system $\omega_{3D}(\phi,s)$ (rotation angle) to test the hypothesis of twist-torsion coupling. However, if the axoneme is not bent in a certain region (local curvature is zero), it is impossible to measure the local bending plane orientation. For small curvatures, it is theoretically possible to calculate the local bending plane orientation, but this calculation is extremely sensitive to the noise of the experimental data (localization errors). We anticipate that the higher the local curvature, the more reliably we can measure the orientation of the local bending plane. Therefore, measurements of torsion as well as local bending plane orientation with respect to the laboratory coordinate system are reliable only in regions where the curvature is high. To determine the error of the local torsion measurement and to set a threshold curvature, above which these measurements can be generally trusted, we used a bootstrapping approach. Here, the 53’000 recorded 3D shapes of 17 axonemes (average *z*-values corrected for single axonemes) serve as the population from which we draw a random sample of 53’000 shapes with repetition. Subsequently, we employ the evaluation pipeline, which calculates the average shape and yields a torsion map. By repeating this procedure for N = 1’000 times, we obtain N torsion maps, from which we can calculate the average torsion map (Figure S3B). Similarly, we obtain the average curvature map from bootstrapping (Figure S3A). From the N torsion maps obtained through bootstrapping, we can calculate the standard deviation for each point in the average torsion map, which is as a proxy for the standard error of the mean (SEM). This allows us to estimate an error for each measured torsion value, resulting in a map of the SEM of torsion (Figure 3F, Figure S3C). A correlation between the curvature map (Figure S3A) and the map of the SEM of torsion (Figure S3C) is clearly visible, which is in support of our method that uses a curvature threshold to define the region-of-trust. Plotting the average torsion values (Figure S3B) as function of the local absolute curvature (absolute values of Figure S3A) yields a plot that shows a large variability of torsion with extremely high values (|$\tau$| > 45 °/µm) at low curvature, which further confirms that torsion is ill defined for low absolute curvature values (Figure S3D). We analyze the curvature-dependent variability of torsion in terms of the interquartile range (IQR) and the standard deviation (SD) as function of absolute curvature (Figure S3E, bin width Δ|*к*| = 0.1 rad/μm). We find that both of these measures of variability steeply decrease with higher absolute curvature before |*к*| = 0.35 rad/μm. The IQR remains approximately constant while the SD further decreases slowly for higher curvature values. Thus, we set a conservative threshold curvature of |*к*| = 0.4 rad/μm, above which the variability in torsion saturates. The calculated SEM of torsion is a function of curvature too and shows the expected inverse relationship (Figure S3F). Here, higher absolute curvature corresponds to smaller SEM of torsion. The SEM of torsion above the threshold curvature |*к*| = 0.4 rad/μm is generally smaller than 4.2 °/µm.

Since the calculation of the orientation angle $\omega_{3D}(\phi,s)$ also depends on the determination of the local bending plane, its measurement error is affected by local curvature in the same way as for the torsion measurement. Thus, we use the same threshold curvature |*к*| = 0.4 rad/μm for the evaluation of $\omega_{3D}(\phi,s)$.

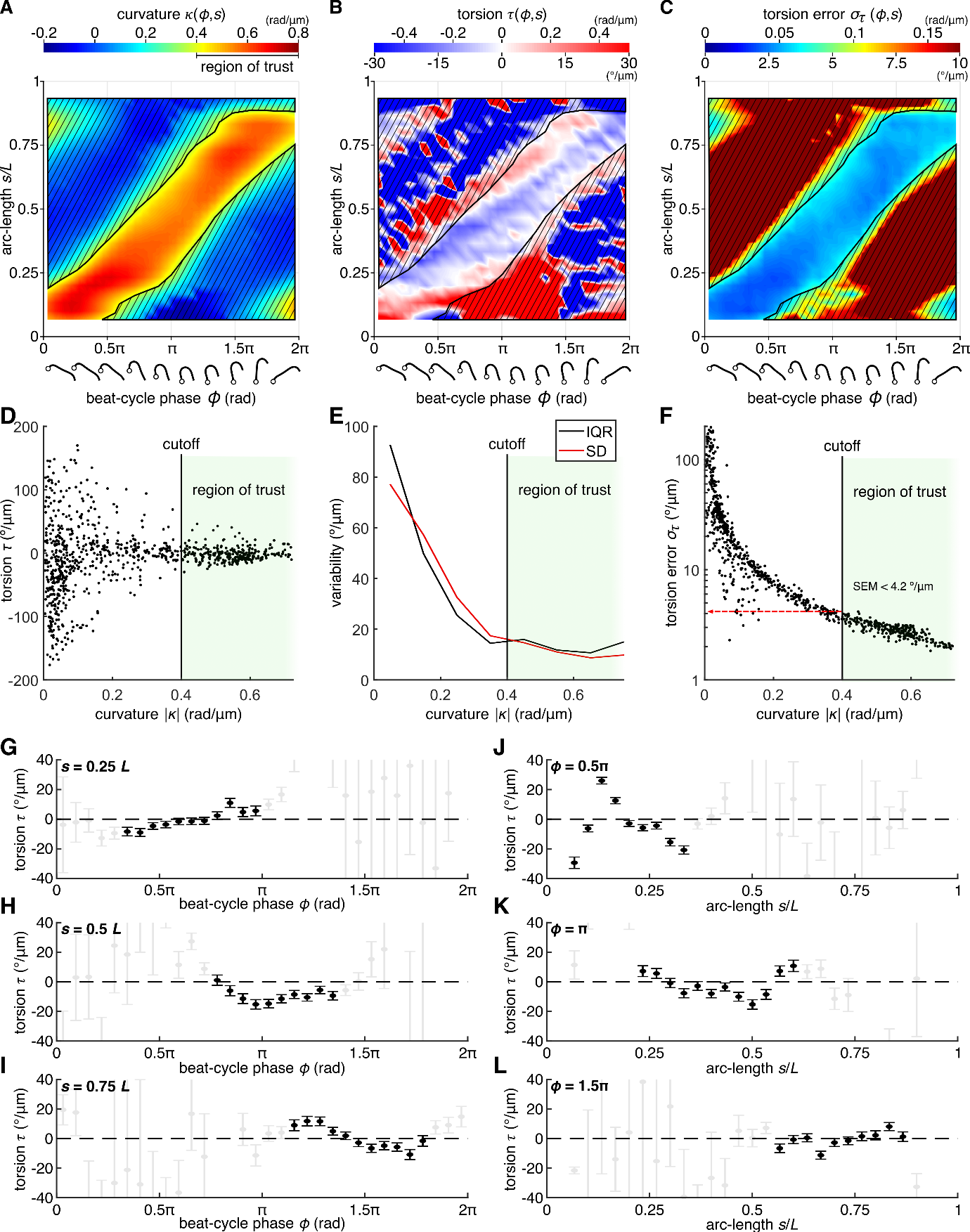

**Figure S3: Torsion error and region-of-trust. (A)** Map of curvature $\kappa$ as function of arc-length *s* and beat-cycle phase $\phi$, calculated as average of N = 1’000 curvature maps that were obtained through bootstrapping. Black lines indicate borders of the region-of-trust ($|\kappa|$> 0.4 rad/µm). **(B)** Map of torsion $\tau$ as function of arc-length *s* and beat-cycle phase $\phi$, calculated as average of N = 1’000 torsion maps that were obtained through bootstrapping. Black lines indicate borders of the region-of-trust ($|\kappa$| > 0.4 rad/µm). **(C)** Map of standard error of mean of torsion $\sigma_{\tau}$ as function of arc-length s and beat-cycle phase $\phi$, calculated as standard deviation of N = 1’000 torsion maps that were obtained through bootstrapping. Black lines indicate borders of the region-of-trust ($|\kappa|$ > 0.4 rad/µm). **(D)** Scatterplot of all torsion values from panel B as function of the absolute local curvature from panel A. The black line marks the threshold curvature for the region-of-trust ($|\kappa|$ > 0.4 rad/µm). **(E)** Analysis of curvature-dependent variability of measured torsion from panel D, depicted as plot of the inter-quartile range (IQR) and the standard deviation (SD) as function of absolute local curvature (bin width = 0.1 rad/µm). The black line marks the threshold curvature for the region-of-trust ($|\kappa|$ > 0.4 rad/µm). **(F)** Scatterplot plot of $\sigma_{\tau}$ from panel C as function of the absolute local curvature from panel A. The black line marks the threshold curvature for the region-of-trust ($|\kappa|$ > 0.4 rad/µm). Torsion errors in the region-of-trust are generally smaller than 4.2 °/µm. **(G-I)** Torsion as function of beat-cycle phase with SEM for *s* = 0.25 *L*, *s* = 0.5 *L*, *s* = 0.75 *L*. Datapoints inside and outside the region-of-trust are black and gray respectively. Datapoints that exceed the axis scaling are not depicted. **(J-L)** Torsion as function of arc-length with SEM for $\phi$ = 0.5π, $\phi$ = π, $\phi$ = 1.5π. Datapoints inside and outside the region-of-trust are black and gray respectively. Datapoints that exceed the axis scaling are not depicted.

**Calculation of torsion amplitude and bending plane rotation amplitude**

The local torsion at a given arc-length position changes during the beat-cycle, which is an indication for dynamic torsion. The change of torsion within our region-of-trust is larger than the estimated measurement error Figure S3G-I). As a lower bound for the peak-to-peak amplitude of dynamic changes in local torsion, we calculate the range of variance within the region-of-trust. The arc-length average of these amplitudes was 21.9 ± 5.8 °/µm (mean ± SD, n = 24, Figure S4B), where amplitudes values above 45 °/µm were neglected.

To test whether the orientation of the bending plane is coupled to the orientation of the axonemal cross-section (Figure 1C, D), we measure the local bending plane orientation with respect to the laboratory coordinate system (orientation angle $\omega_{3D}$, Figure 2C). The resulting map of $\omega_{3D}(\phi,s)$ shows how the orientation of the bending plane changes as function of arc-length and beat-cycle phase (Figure S4A). Similar to the torsion measurement, we conservatively consider values within the region-of-trust only ($|\kappa|$ > 0.4 rad). Within this region-of-trust, we measure the range of variation of the orientation angle $\omega_{3D}(\phi,s)$ to estimate the peak-to-peak amplitude ${\Delta\omega}_{3D}(s)$ as function of arc-length (Figure S4C, Figure 4A). The arc-length average of these amplitudes equals 19.3 ± 7.9 ° (mean ± SD, n = 26). If twist and torsion were coupled (Figure 1E), then the peak-to-peak amplitude ${\Delta\omega}_{3D}(s)$ should be equal to the peak-to-peak amplitude of the cross-section rotation ${\Delta\omega}_{\mathrm{GNP}}(s)$. Since we can only measure ${\Delta\omega}_{3D}(s)$ within the region-of-trust, we compute the peak-to-peak amplitude ${\Delta\omega}_{\mathrm{GNP}}(s)$ within the same region-of-trust for the direct comparison in Figure 4B.

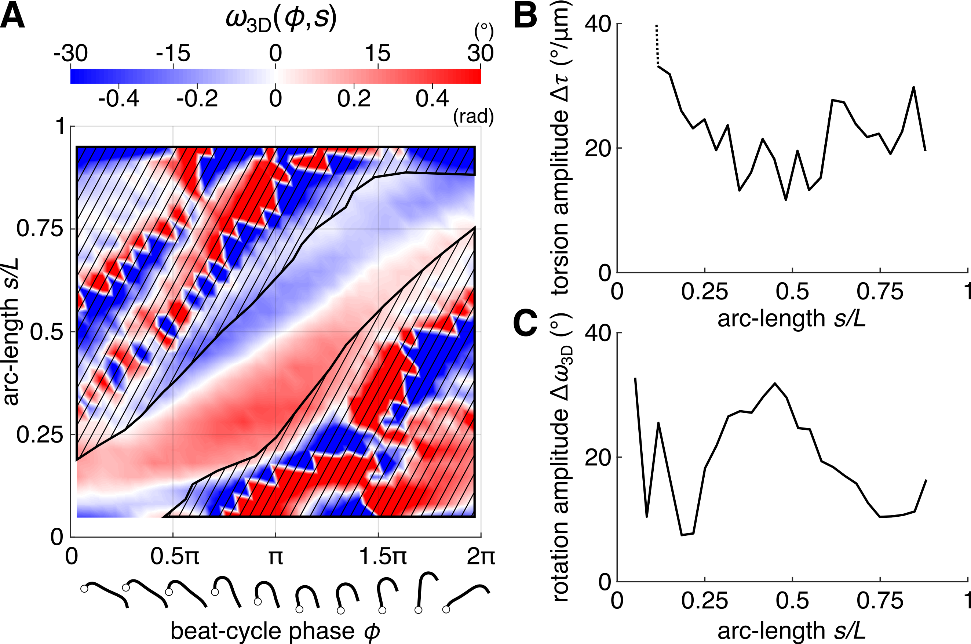

**Figure S4: Bending plane rotation and torsion amplitude. (A)** Map of bending plane orientation with respect to the laboratory coordinate system $\omega_{3D}\left( \phi,s \right)$ as function of beat-cycle phase and arc-length. **(B)** Peak-to-peak torsion amplitudes $\Delta\tau(s)$as function of arc-length s (measured within the region-of-trust). The arc-length average equals 21.9 ± 5.8 °/µm (mean ± SD, n = 24) **(C)** Peak-to-peak bending plane rotation amplitudes $\Delta\omega_{3D}\left( s \right)$as function of arc-length (measured within the region-of-trust). The arc-length average equals 19.3 ± 7.9 ° (mean ± SD, n = 26).

**Measurement of GNP-displacement**

We measure the precise distance between gold nanoparticle (GNP) center and axoneme centerline *d*_C_ in each recorded image of a beating axoneme with attached GNP. Here, we use FIESTA (Ruhnow, Zwicker, and Diez 2011), to first roughly measure the position of the axoneme-bound GNP in the image, in order to select a square shaped region-of-interest (ROI, dimensions 25x25 pixels, Figure S5B,C), which contains the GNP close to its center. To the pixel intensities in this ROI, we then fit a computational model of pixel intensities, which comprises the linear sum of a symmetric 2D Gaussian function $I_{\mathrm{GNP}}(x,y)$ for the GNP and a Gaussian wall along a quadratic spline that locally approximate the axonemal centerline $I_{\mathrm{axo}}(x,y)$ for the axoneme (Figure S5A)

$$I_{\mathrm{GNP}}(x,y)=A_{\mathrm{GNP}}e^{\frac{-{(x-x_{0,\mathrm{GNP}})}^{2}-{(y-y_{0,\mathrm{GNP}})}^{2}}{W_{\mathrm{GNP}}}}$$

$${I_{\mathrm{axo}}\left( x,y \right)=A_{\mathrm{axo}}e}^{\frac{-\left( \begin{matrix} \left( x_{0,\mathrm{axo}}-\alpha\sin\left( \gamma\right) - x \right) \sin\left( \gamma\right)+ \left( y_{0,\mathrm{axo}}-\alpha\cos\left( \gamma\right) - y \right) \cos\left( \gamma\right)+ \\ g \left( \left( x_{0,\mathrm{axo}}-\alpha\sin\left( \gamma\right) - x \right) \cos\left( \gamma\right)+ \left( y_{0,\mathrm{axo}}-\alpha\cos\left( \gamma\right) - y \right) \sin\left( \gamma\right) \right)^{2} \end{matrix} \right)^{2}}{W_{\mathrm{axo}}}}$$

For detailed descriptions of the fit parameters, we refer to Table 1. From the fitting, we obtain the 2D center position of the GNP as well as the 2D centerline of the axoneme within the ROI (Figure S5D), which we use to calculate the normal distance between them, called distance to centerline *d*_C_. Note that we assign a sign to *d*_C_ with reference to the proximal-distal polarity of the axoneme. The sign of *d*_C_ is positive if the angle from the local tangent vector of the axoneme, which points along the axoneme centerline in distal direction, and the vector that points from the centerline position towards the GNP position, measured in counter-clockwise rotation sense, is smaller than π. The measurement of signed distance *d*_C_ in each image enables us to follow the dynamics of GNP displacement relative to the axoneme centerline in subsequent images.

**Table 1: Fit parameters for the measurement of the 2D GNP and axoneme position.** The parameters for the intensity model used for the fit are listed in this table. Through model-fitting, we measure the 2D center position of the GNP as well as the 2D centerline position within the ROI. Here, the 2D centerline shape is approximated by a quadratic spline. We use the 2D centerline position, the 2D GNP center position, and the proximal-distal polarity of the axoneme to determine the signed distance to centerline *d*_C_. A visualization of the model is displayed in Figure S5A.

| sub-model | parameter | description |
| --- | --- | --- |
| $I_{\mathrm{GNP}}$ | $A_{\mathrm{GNP}}$ | amplitude of the GNP signal |
|  | $w_{\mathrm{GNP}}$ | related to square of full-width-at-half-maximum of the GNP signal |
|  | $x_{0,\mathrm{GNP}}, y_{0,\mathrm{GNP}}$ | 2D center position of the GNP |
| $I_{\mathrm{axo}}$ | $A_{\mathrm{axo}}$ | amplitude of axoneme signal |
|  | $w_{\mathrm{axo}}$ | related to square of full-width-at-half-maximum of axoneme signal |
|  | $x_{0,\mathrm{axo}}, y_{0,\mathrm{axo}}$ | reference point in ROI |
|  | $\alpha$ | distance of reference point from vertex of centerline quadratic spline (not strictly needed, but improves robustness of fitting) |
|  | $\gamma$ | angular orientation of the local tangent of the centerline |
|  | *g* | proxy for local curvature of centerline, used in local quadratic spline |

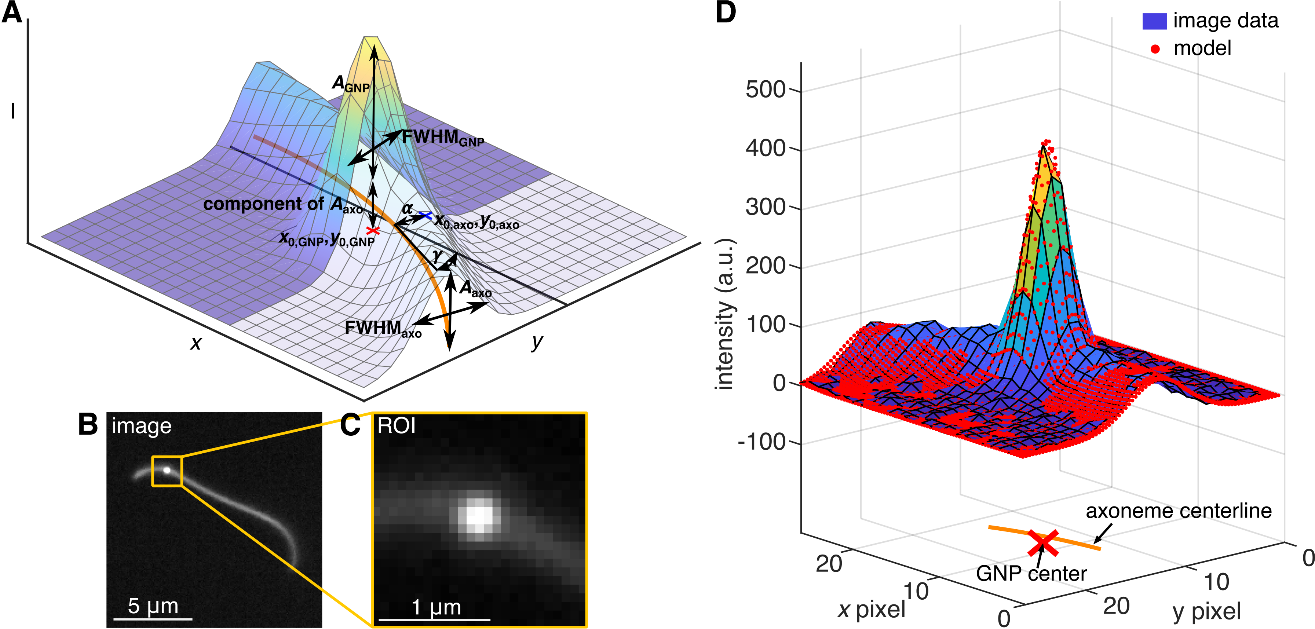

**Figure S5: Fitting of computational model for *d*_C_ measurement. (A)** Visualization of the computational model used to determine the 2D position of the axoneme centerline and the GNP center position. The model is the sum of a Gaussian wall along a quadratic spline (axoneme) and a symmetric 2D Gaussian function (GNP). Table S1 lists all fit parameters. **(B)** Recorded darkfield microscopy image of beating axoneme with attached GNP. Yellow box indicates ROI. **(C)** Zoom into ROI as indicated in panel B. **(D)** Surface plot of the image intensities in the ROI from panel C (background subtracted) with fitted model (red points) and resulting axoneme centerline position (orange line) and GNP center-position (red cross).

**Error estimations of the *d*_C_ measurement**

In the following, we discuss errors of the *d*_C_ measurement, including a curvature-dependent error resulting from model-fitting in our algorithm, two different local tangent angle-dependent errors introduced by both by the algorithm and the imaging system, and finally technical errors with no apparent systematic dependence on dynamic shape and axoneme orientation. We then present a correction method that mitigates local tangent angle-dependent errors. Note that this correction does not affect the reported beat-cycle average *d*_C_($\phi$) but reduces its variance.

**Estimation of the *d*_C_-error related to model-fitting based on simulated imaging data.**

We measure the normal distance *d*_C_ between the (projected) axoneme centerline and GNP center by fitting a combined model function to the intensity distributions in the image as detailed in the preceding section. To determine the existence and extent of bias in the *d*_C_ measurement that may originate from this fitting, we use simulated image data where the ground truth is known. We simulate 2D shapes of the axoneme beat-cycle with bending dynamics described by the prototypical waveform:

$$\kappa\left( s,\phi\right)=A_{\kappa}\sin\left( \phi-\frac{2\pi s}{\lambda}-\frac{\pi}{2} \right)+\kappa_{0}$$

As parameters, we set amplitude $A_{\kappa}$ = 0.5 rad/µm, static curvature $\kappa_{0}$ = 0.3 rad/µm, and wavelength $\lambda$ = *L* = 12 µm. The chosen values for amplitude and static curvature are slightly larger than the corresponding measured values, where we measure an arc-length average amplitude of $A_{\kappa}$ = 0.33 ± 0.06 rad/µm (mean ± SD, n = 27) and an arc-length average static curvature of $\kappa_{0,\exp}$= 0.23 ± 0.06 rad/µm (mean ± SD, n = 27). By choosing slightly larger values for the synthetic data, we aim to rigorously test the performance of our algorithm and to derive a conservative estimate for a curvature-dependent *d*_C_-error. We compute 80 shapes throughout the beat-cycle, each with 328 equidistant datapoints along the arc-length (with $\Delta\phi$ = $\frac{\pi}{40}$ and $\Delta s$ = $\frac{pixel size}{2}$, where the pixel size was 73.33 nm). We then calculate the center position of a hypothetical GNP attached at *s* = *L*/2 and with constant *d*_C_ (ranging between ‑100 nm and 100 nm). From the computed axoneme shape and known GNP positions, we create a simulated image with pixel size 73.33 nm by adding symmetric 2D Gaussian functions to a black image. For the axoneme signal, we add symmetric 2D Gaussian functions with FWHM = 550 nm at each of the 328 datapoints along the arc-length and we set the amplitude of the resulting Gaussian wall equal to 76 (amplitude and FWHM comparable to experimental data). Subsequently, we add one symmetric 2D Gaussian function with amplitude = 279 and FWHM = 550 nm (comparable to experimental data) centered at the known position of the virtual GNP to the image. To mimic experimental image noise, we add shot noise to each pixel (random values drawn independently from a normal distribution with mean µ = 0, standard deviation σ = $\sqrt{pixel intensity}$ ) as well as background noise (random values drawn independently from a normal distribution with mean µ = 47, standard deviation σ = 3.2), comparable to background noise in experimental data. To test for orientation-dependent artifacts as observed in the experimental data, we generate multiple images for each simulated axoneme with virtually attached GNP, where we sampled 64 different orientations of axoneme with GNP in the field of view (each offset by $\frac{\pi}{32}$). To exclude artifacts from pixilation, we subsequently translate axoneme and GNP position together in *x*- and *y*- direction by a random value drawn from normal distributions (mean µ = 0, standard deviation σ = 1 pixel size = 73.33 nm).

We analyzed this synthetic data in exactly the same way as the experimental data: we use FIESTA to approximately trace the position of the GNP in the image. Subsequently we use the evaluation algorithm that we also use to analyze the experimental data (fitting of image intensities), which measures *d*_C_ and we calculate the local 2D curvature $\kappa_{\mathrm{GNP}}$ at the GNP attachment point from the 2D centerline shape in the ROI. We evaluate the performance of the algorithm by calculating the *d*_C_-error $\delta$*d*_C_ as the difference between measured *d*_C_ and true *d*_C_. We observe a correlation of $\delta$*d*_C_ and local $\kappa_{\mathrm{GNP}}$, which depends on the true *d*_C_ (Figure S6E). Since local $\kappa_{\mathrm{GNP}}$ is a function of the beat-cycle (Figure S6D), a curvature-dependent error $\delta$*d*_C_ results in a systematic bias that cannot be averaged out through beat-cycle averaging. This bias is strongest if the particle is attached at *d*_C_ = 100 nm. Here, the curvature dependent $\delta$*d*_C_ could result in a peak-to-peak amplitude of the average *d*_C_ throughout the beat-cycle of ≈ 10 nm for the largest $\kappa_{\mathrm{GNP}}$ amplitudes. For GNPs attached at between *d*_C_ = -100 nm and *d*_C_ = 0 nm, this amplitude is approximately 5 nm. Since these errors are small compared to the average measured peak-to-peak amplitude of approximately 51.0 nm, we conclude that this curvature-dependent *d*_C_-error represents a systematic, but comparatively small artifact.

Using our simulated data, we further show that $\delta$*d*_C_ is correlated with the orientation of the virtual axoneme relative to the coordinate axes (Figure S6F). We observe an orientation-dependent error displaying 4 periods throughout a full rotation of axoneme orientation in the 2D images. This periodicity is likely caused by pixilation, resulting in positive $\delta$*d*_C_ when the axoneme is aligned with the pixel grid (horizontal or vertical) and showing negative $\delta$*d*_C_ when the orientation angle of the axoneme relative to the grid is approximately π/4. The peak-to-peak amplitude is smaller than 5 nm for all values of true *d*_C_ tested. Although these peak-to-peak amplitudes are small, we correct for this orientation-dependent error as shown in the paragraph below.

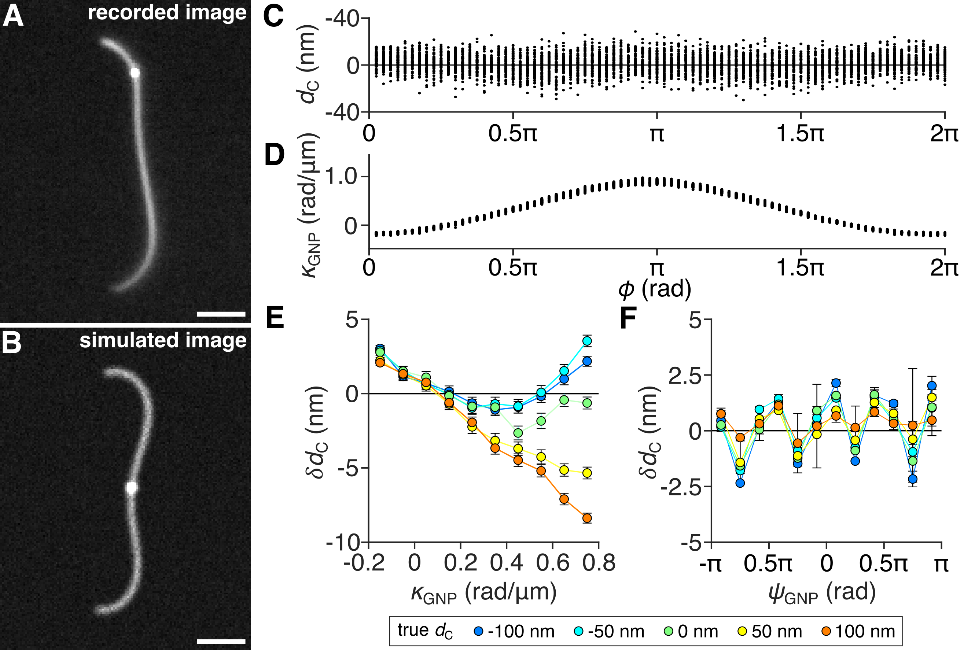

**Figure S6: Validation of the *d*_C_ measurement through model fitting with simulated imaging data. (A)** Recorded darkfield microscopy image of beating axoneme with attached GNP. Scale bar 2 µm**. (B)** Simulated darkfield microscopy image of beating axoneme with attached GNP with known ground truth (known *d*_C_). Scale bar 2 µm. **(C)** Example for measured *d*_C_ from simulated images as function of the beat-cycle phase, black line shows ground truth (known *d*_C_ = 0 nm). **(D)** Example for measured local $\kappa_{\mathrm{GNP}}$ at GNP attachment point in simulated images as function of the beat-cycle phase. **(E)** Average error in *d*_C_-measurement $\delta$*d*_C_ ± SEM as function of local $\kappa_{\mathrm{GNP}}$ (bin width $\Delta\kappa_{\mathrm{GNP}}$ = 0.1 rad/µm for computing averages). Error $\delta$*d*_C_ is calculated as the difference between measured *d*_C_ and known *d*_C_. **(F)** Average error in *d*_C_-measurement $\delta$*d*_C_ ± SEM as function of local tangent angle $\psi_{\mathrm{GNP}}$ (bin width $\Delta\psi_{\mathrm{GNP}}$ = π/6 for computing averages).

**Orientation-dependent *d*_C_-error generated by an imaging artifact**

The measured raw *d*_C_-time profiles from experimental data show features that can be consistently interpreted as a superposition of oscillations with different frequencies (Figure S7E). Using a discrete Fourier transform, we obtain the amplitude spectral density (ASD, which is the absolute of the Fourier transform fft), where we consistently find major peaks at *f*_1_ = beating frequency, *f*_2_ = revolution frequency (2D rotation in FOV), as well as satellite peaks close to integer superpositions *n* *f*_1_ + *m* *f*_2_ (see ASD in Figure S7E). Note that the orientation-dependent *d*_C_-error due to pixilation discussed above for synthetic data (Figure S6F) corresponds to the frequency 4 *f*_2_. For the experimental data, we additionally observe an even more pronounced peak at *f*_2_, which is likely directly related to our imaging system.

As the beating axonemes swim along a circular trajectory in the field of view, the orientation of the axoneme-segment where the GNP is attached (quantified by the local tangent angle of the 2D-projection of the centerline shape in the FOV) changes with the revolution frequency as (Figure S7D). Additionally, the orientation of this segment is a function of the beat-cycle (Figure S7C, D). Thus, the observed ASD is consistent with a superposition of two oscillations: (i) a beat-cycle dependent oscillation and (ii) an orientation-dependent oscillation. Between datasets of different axonemes with GNPs that were recorded in different imaging sessions throughout 2 years, the direction of the orientation-dependent effect in the field of view was conserved (high *d*_C_ at $\psi_{\mathrm{GNP}}$ = -π, π; low *d*_C_ at $\psi_{\mathrm{GNP}}$ = 0) (Figure S7B). By imaging surface-bound axonemes with attached GNPs in different orientations ($\Delta\psi$ ≈ π/18, 200 images each in 36 orientations (Figure S7A, B) and measuring the average *d*_C_ in each orientation, we determine that the orientation-dependent effect occurs even in absence of axoneme activity (Figure S7C). Thus, we conclude that this effect is intrinsic to the optical system and not related to the axonemal beat.

**Correction of orientation-dependent *d*_C_-error**

We are exclusively interested in changes in *d*_C_ due to the axonemal beat. Below, we determine *d*_C_ as function of the beat-cycle phase, where we calculate the average *d*_C_ in small bins throughout the beat-cycle ($\Delta\phi$ = π/16). Since we image the axonemes for multiple revolutions and thus, all beat-cycle phases occur in various orientations, the orientation-dependent effect is expected to not influence the average *d*_C_ as function of the beat-cycle. Nonetheless, we expect this systematic, orientation-dependence *d*_C_-error to increase the estimated SEM of the average *d*_C_. To remove this source of variance, we subtract the orientation-dependent *d*_C_-error for further analysis. To subtract this orientation-dependent *d*_C_-error for single axonemes, we measure the orientation-angle dependent *d*_C_-error $\delta^{\psi}d_{C}$ as the deviation between the total average *d*_C_ and a conditional average *d*_C_ conditioned for a given axoneme orientation angle$\psi_{\mathrm{GNP}}$ ($\Delta\psi$ = π/16). Thus, we obtain a curve of *d*_C_-error as function of orientation angle (32 datapoints), which we linearly interpolate. Using this curve, we calculate the orientation-dependent error for each recorded image, where the orientation angle can be measured. This allows us to calculate a time series of the orientation-dependent error in *d*_C_ (Figure S7F). The ASD of the isolated orientation-dependent error in *d*_C_ shows a peak at the revolution frequency *f*_2_ as well as satellite peaks close to integer superpositions *n* *f*_1_ + *m* *f*_2_, but not at the beating frequency *f*_1_ itself. This supports the notion that *f*_2_ and the satellite peaks originate from the orientation-dependent *d*_C_-error. We subtract the calculated orientation-dependent *d*_C_-error $\delta^{\psi}d_{C}$ from the measured raw *d*_C_-time profile and obtain a corrected *d*_C_-time profile (Figure S7G). The ASD of the corrected a *d*_C_-time profile shows a single peak at the beating frequency *f*_1_ ≈ 60 Hz, and no peaks at the revolution frequency *f*_2_ or any satellite peaks (Figure S7G, see ASD).

By construction, this correction for the orientation-dependent *d*_C_-error should attenuate both the *d*_C_-error arising from our optical imaging system (frequency *f*_2_) as well as pixilation effects affecting model fitting in our algorithm (frequency 4 *f*_2_).

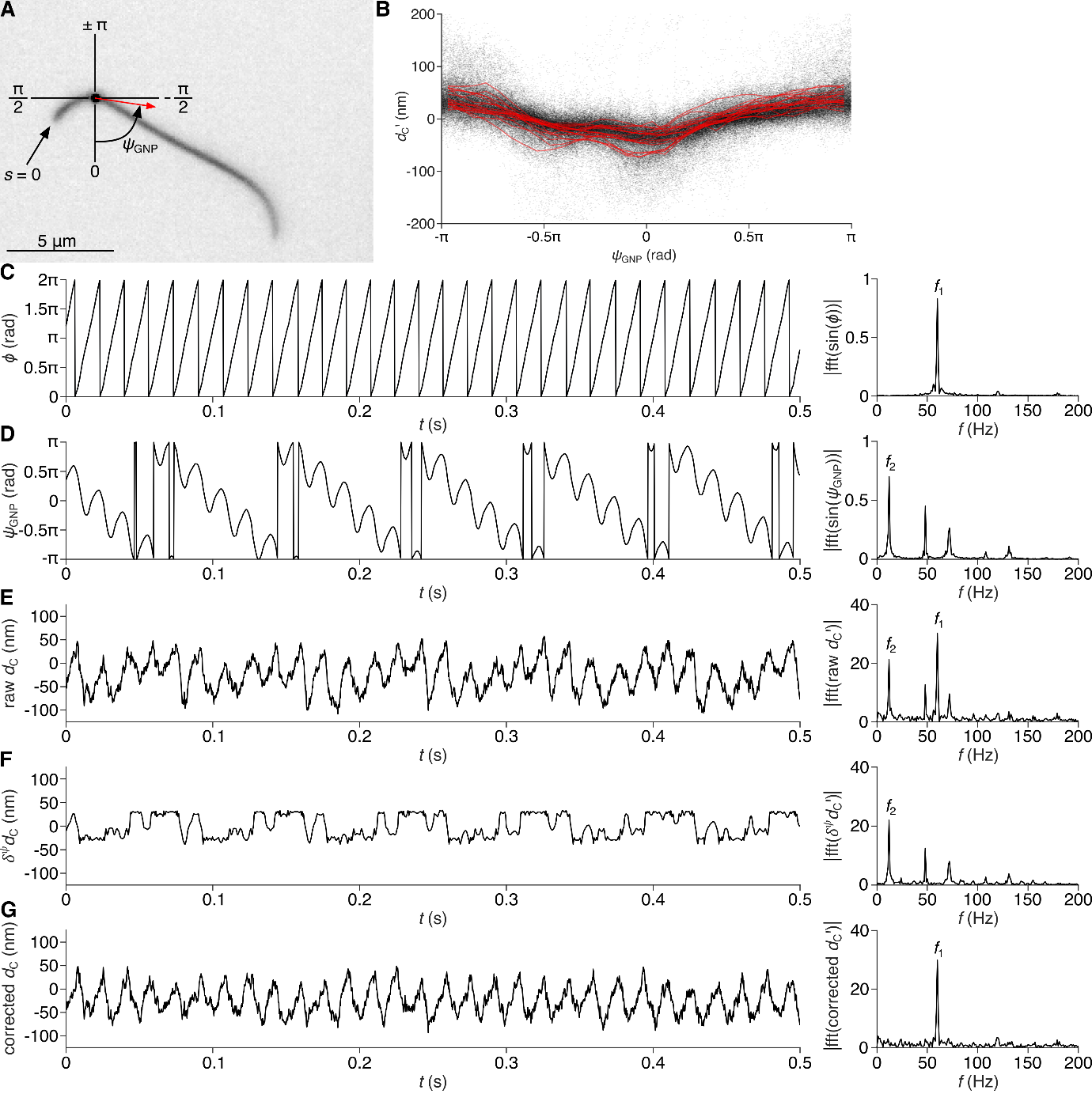

**Figure S7: Correction of orientation-dependent *d*_C_ error. (A)** Reactivated axoneme with attached GNP; red arrow indicates 2D tangent vector at the GNP position. The 2D orientation angle in the field of view of the axoneme at the GNP position$\psi_{\mathrm{GNP}}$ is measured with respect to the coordinate axis as indicated. **(B)** Time-average corrected measured distance to centerline $d_{C}'$ for n = 20 GNPs as function of local 2D tangent angle $\psi_{\mathrm{GNP}}$ in FOV (n = 20 GNPs, total of 150’000 frames). Red curves indicate averages of single axonemes, black dots indicate single datapoints. **(C)** (left) Beat-cycle phase as function of time $\phi(t)$; (right) amplitude spectral density (ASD) of $\sin(\phi(t))$ with a dominant peak at the beating frequency *f*_1_ ≈ 60 Hz. **(D)** (left) Local 2D tangent angle as function of time $\psi_{\mathrm{GNP}}(t)$; (right) ASD of $\sin(\psi_{\mathrm{GNP}}(t)$) with a dominant peak at the 2D revolution frequency *f*_2_ ≈ 12 Hz. **(E)** (left) Measured raw *d*_C_(*t*) as function of time *t* for the GNP that is displayed in panel A; (right) ASD of mean-subtracted raw *d*_C_$'$(*t*) with dominant peaks at the beating frequency *f*_1_ ≈ 60 Hz and at the 2D revolution frequency *f*_2_ ≈ 12 Hz. **(F)** (left) Calculated orientation-dependent *d*_C_-error $\delta^{\psi}d_{C}(t)$ as function of time *t*; (right) ASD of mean-subtracted $\delta^{\psi}d_{C}'$(*t*) with a dominant peak at the 2D revolution frequency *f*_2_ ≈ 12 Hz. **(G)** (left) Corrected *d*_C_(*t*) as function of time *t*; (right) ASD of mean-subtracted corrected *d*_C_$'$(*t*) with a dominant peak at the beating frequency *f*_1_ ≈ 60 Hz.

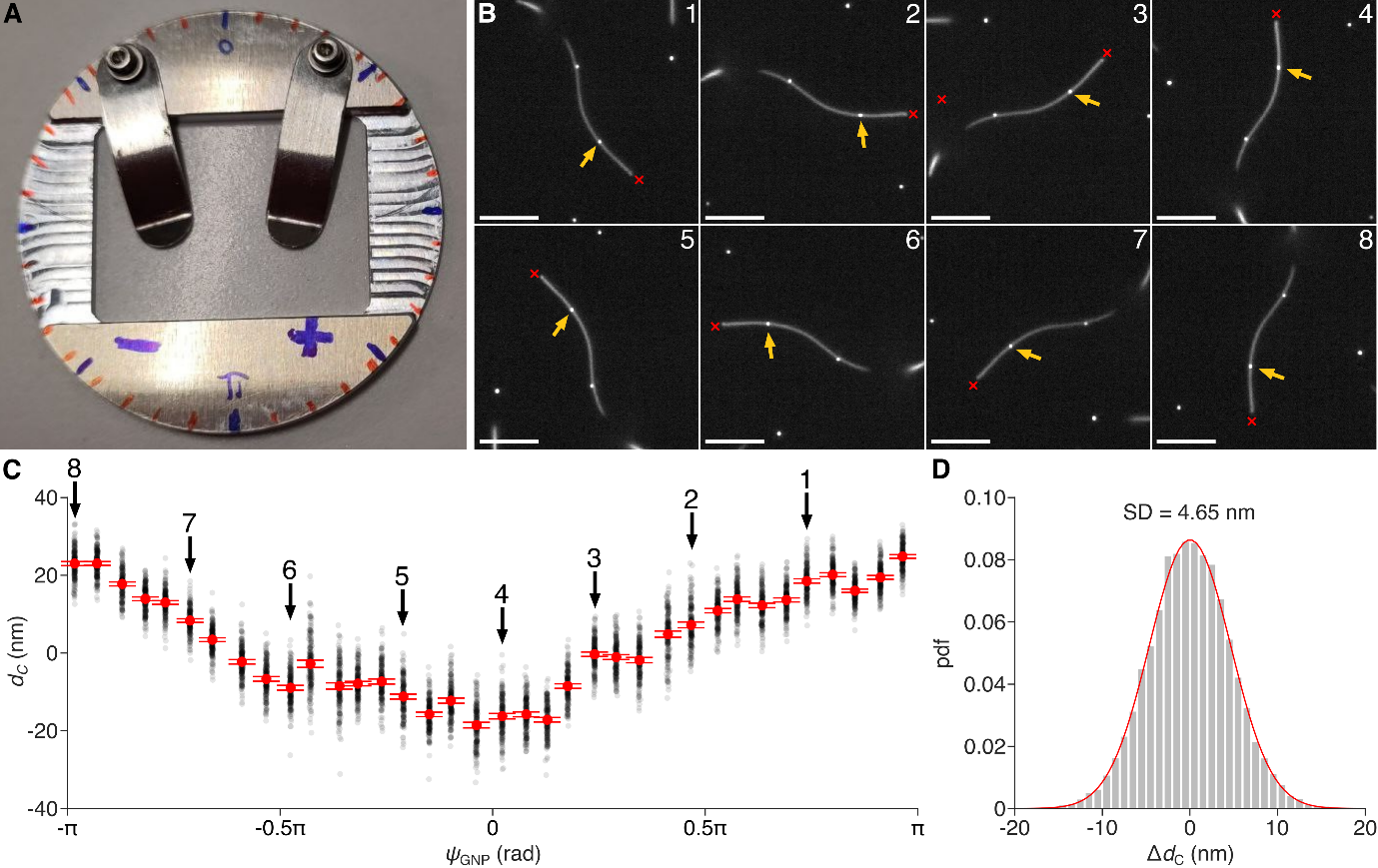

**Figure S8: Measured *d*_C_ changes in orientation-dependent manner in inactive axonemes. (A)** Rotatable stage for imaging samples in different orientations by manual rotation. **(B)** Inactive axoneme with attached GNP imaged in different orientations (red cross: proximal/basal end of the axoneme, yellow arrow: position of GNP, scale bar 5 µm). **(C)** Measured *d*_C_ as function of measured orientation angle $\psi_{\mathrm{GNP}}$ of axoneme section where GNP was attached (immobilized axoneme, no ATP in buffer solution). Displayed are 200 measured datapoints at each sampled orientation with mean ± 95% confidence interval shown in red. Numbers refer to typical example axoneme shown in panel B. **(D)** Typical example of distribution of *d*_C_ deviation around average value for all recorded orientations (n = 7’200 datapoints in total). Fit with normal distribution yields a standard deviation SD of 4.65 nm (R^2^ ≈ 0.998).

**Estimation of random *d*_C_-error in imaging data**

To estimate the random *d*_C_-error, related to image noise, we record representative surface bound axonemes with attached GNPs using the same imaging configuration as for beating axonemes (5’000 fps). To determine the random *d*_C_-error, we used the same images as for measuring orientation dependent *d*_C_-error (Figure S8A-C). However, here we calculate the residuals between the measured *d*_C_ and the average *d*_C_ at a given orientation and subsequently pool these residuals from all orientations. For a typical GNP, the distribution of these residuals is well described by a normal distribution with a standard deviation of 4.65 nm (R^2^ ≈ 0.998, Figure S8D). Thus, we conclude that the random *d*_C_-error in the experimental data, related to image noise, is on the order of 4.65 nm.

**Beat-cycle averaging of *d*_C_ and calculation of the cross-section rotation**

We measure the beat-cycle-dependent rotation of the axoneme cross-section relative to the laboratory frame by evaluating the displacement of GNPs attached to beating axonemes. By measuring the normal distance *d*_C_ between the (projected) axoneme centerline and GNP center, we deduce the circumferential position of the GNP using the known radius of the axoneme (100 nm) and the radius of the GNP (25 nm). As we analyze a 2D projection, this circumferential position is only determined up to a reflection at plane parallel to the image plane. From each image of an axoneme with attached GNP, we measure *d*_C_ as well as the 2D shape of the axoneme to determine the beat-cycle phase $\phi$ (Figure S2A-D). For each axoneme, we superimpose all measurements of *d*_C_ over the beat-cycle (Figure S9). We calculate the average *d*_C_($\phi$) as well as the 95% confidence intervals in 32 bins throughout the beat-cycle ($\Delta\phi$ = π/16, Figure S9: red line is beat-cycle average, width of red line is equal to 95% confidence interval). Due to the high number of measurements (n between 5’000 and 10’000 per single GNP), we can report a significant change in *d*_C_ throughout the beat-cycle for all analyzed axoneme examples. To reliably quantify local cross-section rotation from a *d*_C_-measurement, the GNP should ideally be attached in a circumferential position above or below the axoneme centerline relative to the *z*-axis of the laboratory frame, so that its projected position appears close to the 2D centerline in the 2D image. In contrast, tracking the axonemal cross-section rotation is very unreliable for GNPs that approach the side of the axoneme in the 2D image during the beat cycle (i.e., *d*_C_ ≈ *d*_0_ = 125 nm). This is because once a GNP reaches the side of the axoneme, a change in circumferential position of the GNP only translates into a small change of *d*_C_ in the 2D projection. Because the measured *d*_C_ amplitudes are then small (e.g., on the order of the measurement noise), we only evaluate GNPs where for every $\phi$-bin the 95% confidence intervals of the corresponding *d*_C_($\phi$) lie within an interval of ±125 nm with respect to the 2D centerline. With this rigorous criterium, we exclude GNP measurements that may result in an unreliable measurement of the beat-cycle-dependent rotation of the axonemal cross-section. We note that the majority of GNPs was found attached to the side of axonemes, which strongly limited the number of available samples for this analysis.

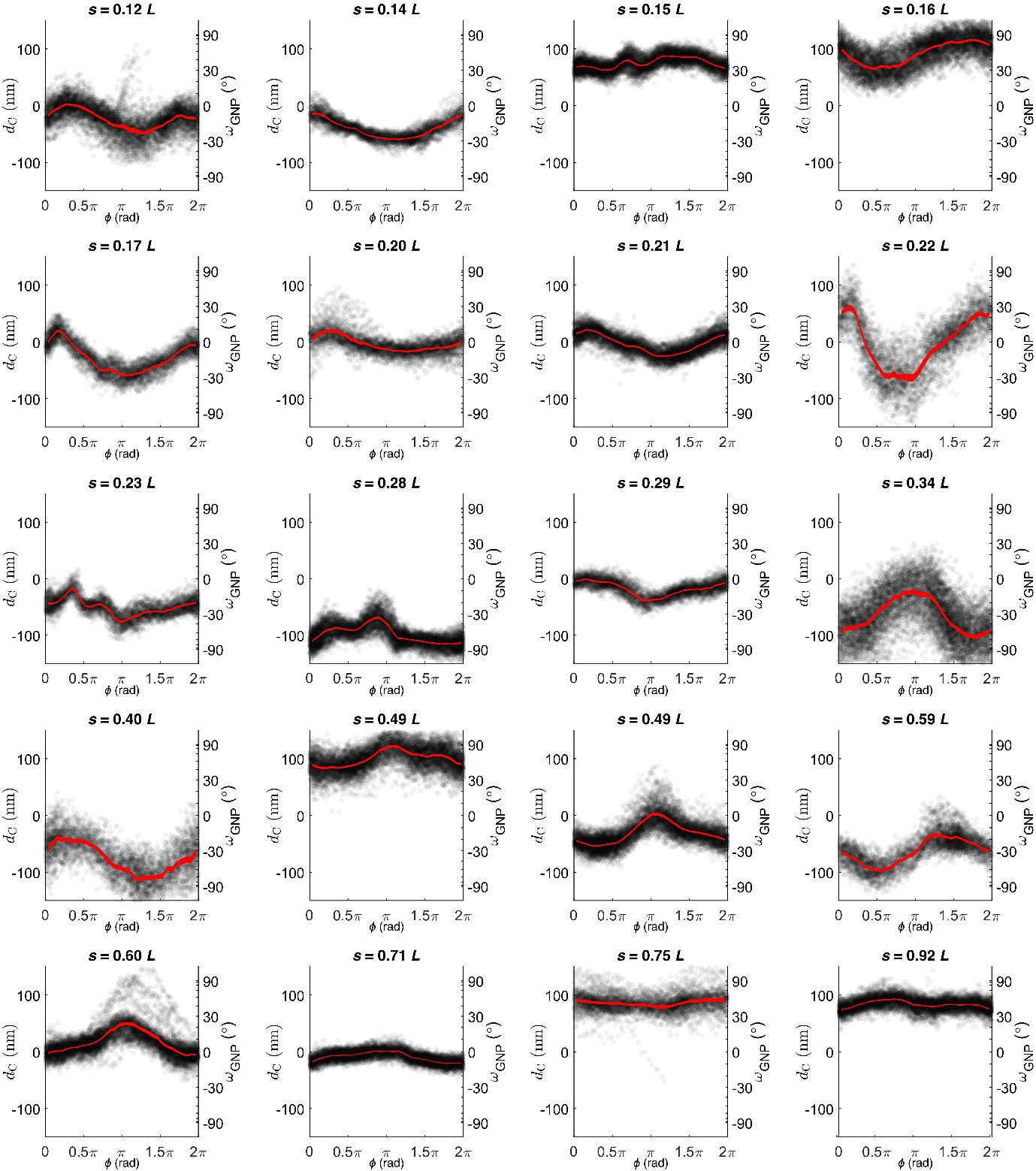

**Figure S9: Measurement of local cross-section orientation.** Each plot shows the corrected *d*_C_ of single recorded images (black dots) and the averaged *d*_C_ (red line, line width = 95% confidence interval, bin width $\Delta\phi=\frac{\pi}{16}$) as function of the beat-cycle phase $\phi$ (horizontal axis). Additionally, the right vertical axis displays the corresponding circumferential position angle $\omega_{\mathrm{GNP}}$ as observed from the laboratory coordinate system (see also Figure 4C). The number of single datapoints in each plot varies between 5’000 and 10’000.

**Map of** $\boldsymbol{\omega}_{\mathbf{GNP}}$ **throughout the beat-cycle and along arc-length**

We measure the beat-cycle-dependent rotation of the local axonemal cross-section relative to the laboratory coordinate system by observing the displacement of GNPs attached to beating axonemes. The peak-to-peak rotation amplitudes ${\Delta\omega}_{\mathrm{GNP}}$ of individual GNPs probing rotations at different arc-length positions are reported in Figure 4E in the main text. In addition, we can combine measurements at different arc-length positions to reconstruct a map of $\omega_{\mathrm{GNP}}$ throughout the beat-cycle and along arc-length. However, since we calculate the angle $\omega_{\mathrm{GNP}}$ from the 2D distance *d*_C_ between the center of the GNP from the axoneme centerline measured in a 2D image as $\omega_{\mathrm{GNP}}$ = cos^-1^ (*d*_C_ / *d*_0_), where *d*_0_ equals 125 nm, $\omega_{\mathrm{GNP}}$ is only determined up to sign. This reflects the fact that from a 2D image, we cannot determine whether the GNP is attached above or below the axoneme relative to the optical axis (Figure S10A). To display $\omega_{\mathrm{GNP}}$ as function of beat-cycle phase $\phi$ and arc-length *s*, we require knowledge about this sign of individually measured $\omega_{\mathrm{GNP}}$. To assign a sign in an unbiased way, we use a purely mechanical argument, which makes the assumption that the cross-section rotation changes continuously in space and time. From the assumption that $\omega_{\mathrm{GNP}}$ changes continuously in time, we obtain a continuous $\omega_{\mathrm{GNP}}$($\phi,s)$ profile as function of beat-cycle phase $\phi$ for fixed arc-length position *s* as shown in Figures 4C and 5C, which, however, is only determined up to a choice of sign. The assumption that $\omega_{\mathrm{GNP}}$ also changes continuously in space allows us to determine this sign. Consider $\omega_{\mathrm{GNP}}(\phi,s_{1})$ and $\omega_{\mathrm{GNP}}(\phi,s_{2})$ for two different GNPs attached at subsequent arc-length positions *s*_1_ and *s*_2_. As these profiles will likely oscillate around different angular positions that reflect different circumferential positions of attachment, we first subtract the mean between the minimum value and the maximum value for each $\omega_{\mathrm{GNP}}$ oscillation. Next, we calculate the sum of squared residuals *S*_+_ and *S*_−_ between $\omega_{\mathrm{GNP}}\left( \phi,s_{1} \right)$ and $\omega_{\mathrm{GNP}}\left( \phi,s_{2} \right)$ for both choices of sign for $\omega_{\mathrm{GNP}}\left( \phi,s_{2} \right)$ (Figure S10B, C)

$$S_{\pm}=\sum_{\phi= \Delta\phi}^{2\pi} \left( \omega_{\mathrm{GNP}}\left( \phi,s_{1} \right)\mp\omega_{\mathrm{GNP}}\left( \phi,s_{2} \right) \right)^{2}$$

We then select the sign, for which the sum of residuals *S* is smaller. This ensures that the axonemal cross-section rotation changes continuously in space.

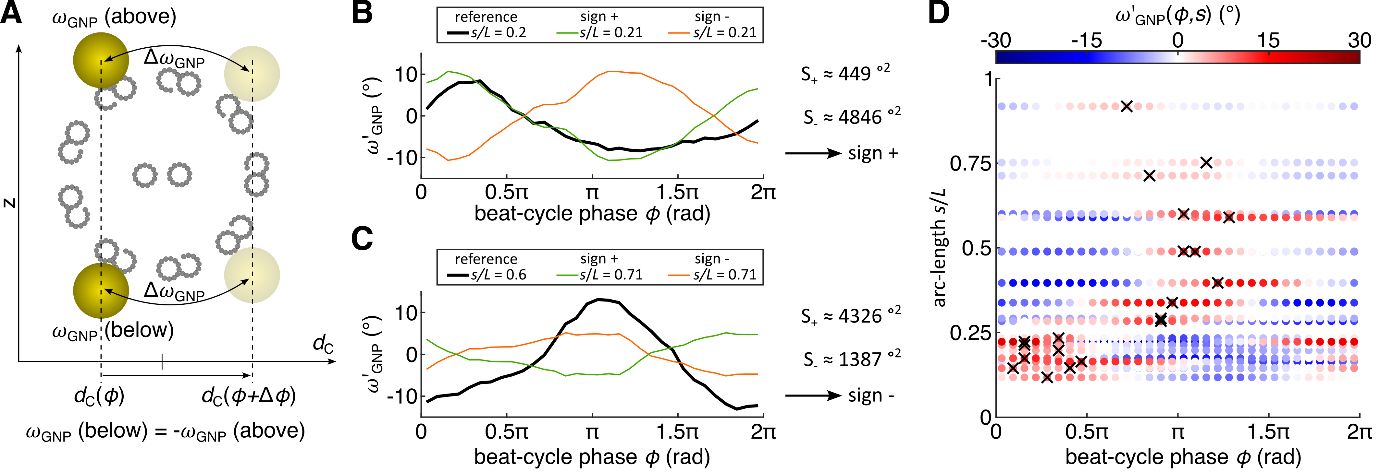

**Figure S10: Map of cross-section rotation relative to the laboratory coordinate system requires choice of sign*.* (A)** Measurement of $\omega_{\mathrm{GNP}}$ from 2D images cannot determine the sign of the local cross-section rotation angle, as we cannot determine whether the GNP is above or below the axoneme relative to the optical axis of the microscope. **(B)** To assign a sign in an unbiased way, we assume that the cross-section rotation changes continuously in space. Here, two rotation angle profiles $\omega_{\mathrm{GNP}}(\phi,s_{1})$ and $\omega_{\mathrm{GNP}}(\phi,s_{2})$ for subsequent arc-length positions *s*_1_ and *s*_2_ are shown (corrected for mean) for different choices of global sign for the second profile at *s*_2_ = 0.21 *L* (green: sign = +1, orange: sign = -1), while the first profile at *s*_1_ = 0.2 *L* serves as reference (black). In this example, the green curve (sign = +1) is closer to the reference, reflected by a value of *S*_+_ smaller than *S*_−_; hence, the sign +1 is chosen. **(C)** Example similar to panel B, but where *S*_+_ is larger than *S*_−_; hence, the red curve corresponding to the sign -1 is chosen. Note that for this example, the two subsequent arc-length positions *s*_1_ = 0.6 *L* and *s*_2_ = 0.71 *L* are further apart, and hence the corresponding rotation angle profiles differ more, making the choice of sign more difficult. **(D)** After iteratively assigning signs to all $\omega_{\mathrm{GNP}}(\phi,s)$, we reconstruct a map of $\omega_{\mathrm{GNP}}(\phi,s)$ whose global sign is still arbitrary, but the signs between individual $\Delta\omega_{\mathrm{GNP}}(\phi)$ are consistent based on the assumption that the cross-section rotation changes continuously in space and time. To outline possible spatio-temporal trends, we mark the maxima of each $\omega_{\mathrm{GNP}}(\phi)$ (black crosses).

**Calculation of the twist-free material frame and estimation of the local cross-section rotation observed from the laboratory coordinate system**

*Coordinate frames for beating axonemes.* We discuss and introduce different frames of reference, to describe the three-dimensional shapes as well as the structure of beating axonemes. Both are partially characterized by the three-dimensional centerline $\mathbf{r}(s)$, which is a curve in three-dimensional space that is parameterized by the arc-length *s* where 0 ≤ *s* ≤ *L* (*L* = contour length of the centerline). We use those reference frames to characterize both the shape and the mechanical deformations of the axoneme with prescribed centerline. In general, a frame can be thought of as the 3D centerline with a “weather-vane” that slides along it, which defines three-unit vectors ($\mathbf{a}_{1}, \mathbf{a}_{2},\mathbf{a}_{3}$) at every position $\mathbf{r}(s)$. Specifically, $a_{3}\left( s \right)$ is equal to the tangent vector of the centerline $\partial_{s}\mathbf{r}\left( s \right)=\mathbf{t}\left( s \right)=\mathbf{a}_{3}(s)$, $\mathbf{a}_{1}\left( s \right)$ points in the direction of the weather-vane and $\mathbf{a}_{2}\left( s \right)= \mathbf{a}_{1}\left( s \right) \times\mathbf{a}_{3}\left( s \right)$ completes the triad of mutually orthogonal vectors.

We use three different frames to describe the shape and the structure in different scenarios:

- the material frame (commonly used in physics and engineering to describe material properties)
- the Frenet-Serret frame (commonly used in mathematics and geometry to describe the shapes of three-dimensional curves)
- the rotation-free normal frame put forward by Bishop and Wiggins (Bishop 1975; Goldstein, Powers, and Wiggins 1998)

*The material frame* characterizes the axoneme as a physical object of non-zero diameter that
can undergo elastic deformations. The material frame at a given arc-length position is best defined in terms of the cross-section of the axoneme at this point. Specifically, at every arc-length position *s*, we define a set of three vectors $\mathbf{e}_{1}, \mathbf{e}_{2},\mathbf{e}_{3}$ (Figure 1A), where

- $\mathbf{e}_{3}\left( s \right)$ is equal to the tangent vector of the centerline $\mathbf{t}\left( s \right)=\mathbf{e}_{3}\left( s \right)$
- $\mathbf{e}_{1}\left( s \right)$ and $\mathbf{e}_{2}\left( s \right)$ span the plane of the cross-section
- $\mathbf{e}_{1}\left( s \right)$ is locked to a specific position in the axoneme cross-section (Figure 1C). For simplicity, we decide to lock $\mathbf{e}_{1}\left( s \right)$ to the inter-doublet bridge, which naturally introduces asymmetry to the cross-section. The unit vector $\mathbf{e}_{1}\left( s \right)$ can be thought of as a weather vane that slides along $\mathbf{r}(s)$ and follows the position of the bridge.
- $\mathbf{e}_{2}\left( s \right)= \mathbf{e}_{1}\left( s \right) \times\mathbf{e}_{3}\left( s \right)$

The material frame is useful to describe deformations of the axoneme. If the axoneme is deformed, the three vectors rotate as function of arc-length with rates of rotation (units: °/μm or rad/μm), which describe elastic deformations

- $\Omega_{1}$= rate of rotation around $\mathbf{e}_{1}$, equal to bending in direction of ± $\mathbf{e}_{2}$
- $\Omega_{2}$= rate of rotation around $\mathbf{e}_{2}$, equal to bending in direction of ± $\mathbf{e}_{1}$
- $\Omega_{3}$= rate of rotation around $\mathbf{e}_{3}$, equal to twist

This can be equivalently expressed in compact form as

$$\partial_{s}\mathbf{e}_{j}=\boldsymbol{\Omega}\times\mathbf{e}_{j} , j=1,2,3$$

(1)

with

$$\boldsymbol{\Omega}=\Omega_{1}e_{1}+ \Omega_{2}e_{2}+\Omega_{3}e_{3}$$

The elastic energy stored in a deformed axoneme is given to leading order according to Cosserat rod theory as

$$E= \int_{0}^{L} ds\frac{k_{1}}{2}\Omega_{1}^{2}+ \frac{k_{2}}{2}\Omega_{2}^{2}+\frac{k_{3}}{2}\Omega_{3}^{2}$$

(2)

Here, $k_{1}$and $k_{2}$ denote anisotropic bending stiffnesses and $k_{3}$ denotes the twist rigidity.

In general, the material frame describes how the physical axoneme deforms to achieve the three-dimensional shape that we measure in our experiments.

*The Frenet-Serret frame* is a purely geometric description of the 3D centerline of the
axoneme (neglecting the fact that the axoneme is a physical object with non-zero diameter). The three vectors of the Frenet-Serret frame at a given centerline point **r**(*s*) are

- $\mathbf{t}\left( s \right)$ is the tangent vector of the centerline, which is equal to $\mathbf{e}_{3}\left( s \right)$
- $\mathbf{n}\left( s \right)$ is the normal vector, which together with $\mathbf{t}\left( s \right)$ spans the local bending plane (also generally referred to as “the osculating plane” of a 3D curve)
- **b**$\left( s \right)$ is the binormal vector, which is perpendicular to the local bending plane.

Three-dimensional shapes of the axonemal centerline are fully described (excluding rigid body transformations such as translations and rotations) by

- $\kappa\left( s \right)$ is the signed three-dimensional curvature of the centerline, which (up to the sign) is defined as the inverse radius of the osculating circle that matches the centerline curve at **r**(*s*) (units: °/μm or rad/μm).
- $\tau\left( s \right)$ is the torsion, which describes the rotation rate of the local bending plane of the centerline (around $t\left( s \right)$) as function of arc-length (units: °/μm or rad/μm).

Strictly, the Frenet-Serret frame is not defined at inflection points of the centerline, where the curvature vanishes. In practice, one may still compute a Frenet-Serret at these inflection points by requiring that the Frenet-Serret frame should change continuously as function of
arc-length *s*. This requirement also allows to define a global sign for the curvature for the entire shape (Jikeli et al. 2015). Computing torsion in regions where curvature is low, however, is sensitive to measurement errors.

Mathematical formulas allow to compute curvature and torsion of the 3D shape from the bending rates of the material frame (two quantities computed from three quantities), but it is not possible to conversely infer these bending rates from curvature and torsion (three independent quantities cannot be inferred from two quantities).

*Arbitrary frames*. The material frame and the Frenet-Serret frame are just two examples of frames of the space curve **r**(*s*). To highlight the general framework, we briefly consider the concept of a general frame with a family of orthonormal unit vectors $\mathbf{a}_{1}(s), \mathbf{a}_{2}(s),\mathbf{a}_{3}(s)$ parameterized by arc-length *s* with $\mathbf{a}_{3}(s)$ = **t**(*s*) and $\mathbf{a}_{2}\left( s \right)= \mathbf{a}_{3}\left( s \right)\times\mathbf{a}_{1}(s)$. Then, equation (1) can be written in general form as $\partial_{s}\mathbf{a}_{j}=\boldsymbol{\Omega}\times\mathbf{a}_{j}$ (*j* = 1,2,3), explicitly

$$\partial_{s}\mathbf{a}_{1}\left( s \right)= + \alpha_{2}\mathbf{a}_{3}-\alpha_{3}\mathbf{a}_{2}$$

$$\partial_{s}\mathbf{a}_{2}\left( s \right)=- \alpha_{1}\mathbf{a}_{3} +\alpha_{3}\mathbf{a}_{1}$$

$$\partial_{s}\mathbf{a}_{3}\left( s \right)=+ \alpha_{1}\mathbf{a}_{2}-\alpha_{2}\mathbf{a}_{1}$$

(3)

For the material frame, we simply have $\mathbf{a}_{j}=\mathbf{e}_{j}$ and $\alpha_{j}=\Omega_{j}$ with *j* = 1,2,3. In this case, the rotation rates $\alpha_{j}$ denote actual rates of elastic deformation. For the Frenet-Serret frame, we have $\mathbf{a}_{1}=\mathbf{n}$, $\mathbf{a}_{2}=\mathbf{b}$, $\mathbf{a}_{3}=\mathbf{t}$, with rotation rates $\alpha_{1}=\kappa$,$\alpha_{2}=0$, $\alpha_{3}=\tau$. Here, rotation rates are given by curvature and torsion, given that $\alpha_{2}$ is identically zero.

*Relations between arbitrary frames*. Any two arbitrary frames $\mathbf{a}_{j}^{'}(s)$, *j*=1,2,3 and $\mathbf{a}_{j}^{''}(s)$, *j* = 1,2,3 are related by local rotations with the arc-length dependent rotation angle $\theta(s)$ as

$$\mathbf{a}_{1}^{''}=\cos\left( \theta\right)\mathbf{a}_{1}^{'}+\sin(\theta)\mathbf{a}_{2}^{'}$$

$$\mathbf{a}_{2}^{''}=-\sin\left( \theta\right)\mathbf{a}_{1}^{'}+\cos(\theta)\mathbf{a}_{2}^{'}$$

$$\mathbf{a}_{3}^{''}=\mathbf{a}_{3}^{'}$$

(4)

This equation can be rewritten in a complex notation as

$$\mathbf{a}^{''}=\mathbf{a}^{'} e^{i\theta}$$

(5)

where, for notational convenience, we introduced the complex vectors $\mathbf{a}^{'}=\mathbf{a}_{1}^{'}+i\mathbf{a}_{2}^{'}$and
$\mathbf{a}^{''}=\mathbf{a}_{1}^{''}+i\mathbf{a}_{2}^{''}$. The rotation rates of the two frames are related by

$$(-i\alpha_{1}^{''}+\alpha_{2}^{''})=\left( -i\alpha_{1}^{'}+\alpha_{2}^{'} \right) e^{i\theta}$$

(6)

where the expressions in parenthesis can be interpreted as complex bending rates, while

$$\alpha_{3}^{''}\left( s \right)=\alpha_{3}^{'}\left( s \right)-\partial_{s}\theta(s)$$

(7)

We emphasize that a global rotation with constant $\theta\left( s \right)\equiv\theta_{0}$ would not change $\alpha_{3}$, but an arc-length-dependent rotation does. Note that the unsigned curvature can be interpreted as the modulus of the complex rotation rate as $\left| \kappa\right|=\left| - i\alpha_{1}+\alpha_{2} \right|$, which by Equation (7) is independent of choice of frame und thus an invariant of the curve.

*The rotation-free normal frame* is a special frame, for which the rotation rate $\Omega_{3}\left( s \right)$ around the tangent is identically zero.

If the rotation-free normal frame is equal to the material frame, this frame describes a scenario where the twist rigidity $k_{3}$is infinite and the 3D shape emerges from bending ($\Omega_{1}$ and $\Omega_{2}$) exclusively ($\Omega_{3}$ = 0) (Figure 1D). This frame has been originally devised by Darboux and Hasimoto (see refs. in (Goldstein, Powers, and Wiggins 1998)), and discussed in (Bishop 1975), where it was called relatively parallel adapted frame. Equation (8) provides a recipe how to construct a rotation-free normal frame. Here, we may start with an arbitrary frame $\mathbf{a}_{j}^{'}(s)$, *j*=1,2,3 with a rotation rate of $\alpha_{3}$(s) and define (Goldstein, Powers, and Wiggins 1998)

$$\theta\left( s \right)= \theta_{0}+\int_{0}^{s} ds \alpha_{3}(s)$$

(8)

where $\theta_{0}$ is an arbitrary constant. Using this frame as a start frame $\mathbf{a}_{i}^{'}=\mathbf{a}_{i}$ and applying a local rotation with this arc-length-dependent rotation angle according to Equation (7) defines a new frame $\mathbf{m}\left( s \right)=\mathbf{a}_{1}^{''}(s)$, $\mathbf{l}\left( s \right)=\mathbf{a}_{2}^{''}(s)$. By construction, this frame is rotation-free, i.e., $\alpha_{3}^{''}\left( s \right) = 0$.

*Special frames as limit cases of the material frame****.*** If only the centerline $\mathbf{r}(s)$ is known, it is impossible to infer the correct material frame $\mathbf{e}_{j}(s)$, j=1,2,3 without further assumptions. If we can assume, however, that the deformed axoneme assumes a minimum of elastic energy as given by Equation (2), subject to the constraint that the centerline $\mathbf{r}(s)$ is kept fixed, we can numerically compute the material frame by non-linear optimization. The result of this optimization takes a particularly simple form in special limit cases

- $k_{1}\gg k_{2},k_{3}$: in this case, $\Omega_{1}\left( s \right)=0,$ i.e. bending of the axoneme is only possible in the $\mathbf{e}_{1}$ direction (Figure 1C), corresponding to an infinitely stiff cross-bridge. In this limit case, the material frame that minimizes the elastic energy is equal to the Frenet-Serret frame $\mathbf{e}_{1}=\mathbf{n}$,$\mathbf{e}_{2}=\mathbf{b}$,$\mathbf{e}_{3}=\mathbf{t}$. In particular, twist equals torsion

$$\Omega_{3}\left( s \right)=\tau$$

(9)

This means that in this scenario, torsion can only occurs only if the axoneme is twisted. In the main text, we term this limit case twist-torsion coupling.

- $k_{3}\gg k_{1},k_{2}$: in this case, the twist rigidity is infinite (Figure 1D), hence $\Omega_{3}\left( s \right)=0$. In this limit case, the material frame that minimizes the elastic energy is given by a twist-free normal frame $\mathbf{e}_{1}=\mathbf{m}$,$\mathbf{e}_{2}=\mathbf{l}$,$\mathbf{e}_{3}=\mathbf{t}$. Twist is zero

$$\Omega_{3}\left( s \right)=0$$

(10)

This means that in this scenario, torsion occurs in the absence of twist, which we call twist-free torsion.

The two discussed scenarios are limit cases. While it is conceivable that the material frame of a real axoneme represents an intermediate between these two limit cases, we focus our interpretation efforts on determining the more likely limit case.

*Computation of the twist-free normal frame*. For the numerical computation of a twist-free normal frame, one may start, e.g., with an empirical estimation of the Frenet-Serret frame. Next, the rotation of this frame can be undone using Equation (8). This procedure is not unique, as the choice of the constant $\theta_{0}$ in Equation (8) is arbitrary, resulting in a family of equivalent normal frames that are related to each other by a global rotation. This non-uniqueness becomes relevant, when a whole beat-cycle with centerline shapes $\mathbf{r}\left( s,\phi\right)$ that depends on the phase $\phi$ of the beat is to be considered (for example, to predict the motion of a GNP rigidly attached to the axoneme), as now a global rotation angle $\theta_{0}(\phi)$ has to be chosen for each phase $\phi$.

We can fix this gauge freedom by a hydrodynamic argument: Helmholtz’ principle of minimal hydrodynamic dissipation (Happel and Brenner 1983) implies that any variation of the geometry of the axoneme by a degree of freedom *q* (including the choice of $\theta_{0}(\phi)$) should minimize the total hydrodynamic dissipation rate $R$, i.e., the conjugate generalized hydrodynamic friction force $P_{q}=\delta R/\delta\dot{q}$ should vanish (Solovev and Friedrich 2021). Ignoring long-range hydrodynamic interactions between axonemal cross-sections, the generalized hydrodynamic friction force *P* conjugate to $\theta_{0}$ is approximated by

$$P_{q}= \zeta\int_{0}^{L} ds \partial_{\phi}\mathbf{e}_{1}\cdot\mathbf{e}_{2}$$

$$= \zeta\int_{0}^{L} ds \left[ \partial_{\phi}\mathbf{a}_{1}\cdot\mathbf{a}_{2}-\partial_{\phi}\theta_{0} \right]$$

(11)

where $\zeta$ is a friction coefficient. Setting *P* = 0 implies

$$\partial_{\phi}\theta_{0}=\frac{1}{L}\int_{0}^{L} ds \partial_{\phi}\mathbf{a}_{1}\cdot\mathbf{a}_{2}$$

(12)

Equation (12) has a simple geometrical interpretation:$\partial_{\phi}\theta_{0}$ should be chosen such that the integrated local rotations of the cross-sections are minimal. Because $\partial_{\phi}\theta_{0}$ should be a 2π-periodic function of $\phi$, we should have $\int_{0}^{2\pi} d\phi\partial_{\phi}\theta_{0}= \theta_{0}\left( 2\pi\right)-\theta_{0}\left( 0 \right)=0$. If we apply Equation (12) to experimental data on a discrete $\left( s,\phi\right)$-grid, possibly subject to measurement noise, this condition will not exactly be fulfilled, prompting a modification of Equation (12). For discrete data with arc-length spacing $\Delta s$ and beat-cycle discretization $\Delta\phi$, the equivalent of Equation (12) reads $\theta_{0}\left( \phi_{j+1} \right)=\theta_{0}\left( \phi_{j} \right)+\Delta\theta_{0,j}$ with rotation angle increments

$$\Delta\theta_{0,j} =\frac{\Delta s}{L} \sum_{k=1}^{N_{s}} \left[ \mathbf{a}_{1}\left( s_{k},\phi_{j}+\Delta\phi\right)- \mathbf{a}_{1}\left( s_{k},\phi_{j} \right) \right]\cdot\mathbf{a}_{2}\left( s_{k},\phi_{j} \right)$$

(13)

where $s_{k}=k\Delta s$ with $N_{s}=L/\Delta s$ being the number of control points along the axonemal arc-length and $\phi_{j}=j \Delta\phi$ with $N_{\phi}=2\pi/\Delta\phi$ being the number of subsequent axonemal shapes in one beat-cycle. We can use least-square optimization to find an approximate solution to the generalized force balance equations $P\left( \phi_{j} \right)=0$ for *j =* 1*,…,*$N_{\phi}$ that ensures the periodicity condition also in the case of discrete data, with explicit solution

$$\theta_{0}\left( \phi_{j+1} \right)= \theta_{0}\left( \phi_{j} \right)+\Delta\theta_{0,j}-\frac{\Delta\phi}{2\pi}\sum_{k=1}^{N_{\phi}} \Delta\theta_{0,k}$$

(14)

This procedure defines an (almost unique) normal frame that only depends on the choice of a single constant $\theta_{0}\left( \phi=0 \right)$ at the beginning of the beat-cycle. This choice, however, does not affect the prediction of the rotation angle $\omega_{no twist}(\phi,s)$ of a GNP attached to an axoneme, whose material frame is given by any of these twist-free normal frames.

Using the procedure described above, we first calculate twist-free normal frames from the Serret-Frenet-frames of the 32 experimentally obtained shapes that comprise the average 3D waveform (using Equation (8)). As detailed above, individual twist-free normal frames have arbitrary global rotation angles (Figure S11A). We align global rotation angles of subsequent beat-cycle phases using Equation (14). This results in 32 twist-free normal frames throughout the beat-cycle (Figure S11A).

If the twist-free scenario (Figure 1D) is assumed to be true, these twist-free normal frames will represent a material frame. Since we measured the 3D shapes and 3D orientation of the axoneme, we can calculate how these twist-free material frames are oriented in 3D as function of the beat-cycle phase. This allows us to predict the movement of a GNP that is attached to such a material frame and calculate the local cross-section rotation relative to the laboratory frame $\omega_{no twist}(\phi,s)$. We calculate a map of $\omega_{no twist}(\phi,s)$ (Figure S11B) and present the $\omega_{no twist}(\phi)$ profiles for all arc-length positions (Figure S11C). This map $\omega_{no twist}(\phi,s)$ shows features that are consistent with a travelling wave of cross-section rotation as observed from the laboratory coordinate system. Note that in this scenario, the material frame of the axoneme remains completely twist-free and the apparent cross-section rotation arises solely from 3D shape changes and resulting changes in 3D orientations with respect to the laboratory coordinate system.

We observe that the peak-to-peak amplitude$\Delta\omega_{no twist}(s)$ as function of arc-length shows values that are smaller than the experimentally measured local cross-section rotation $\Delta\omega_{\mathrm{GNP}}(s)$ (Figure S11, Figure 4E, Figure 5A). We calculate an arc-length average of $\Delta\omega_{no twist}(s)$ of 7.2 ± 0.3 ° (mean ± SEM, n = 28).

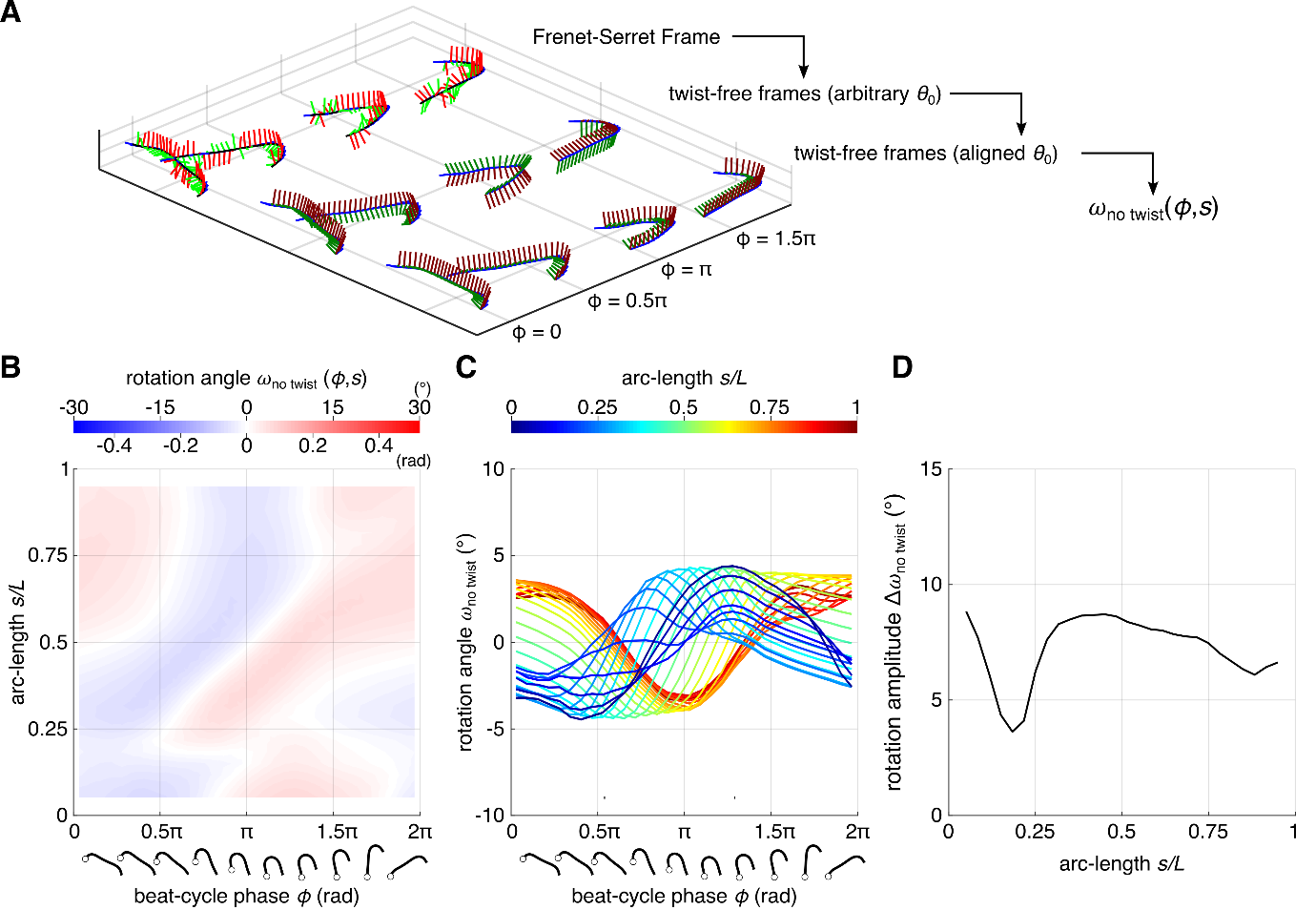

**Figure S11: GNP rotation on a twist-free axoneme. (A)** We calculate the hypothetical twist-free normal material frame from the Frenet-Serret frame (using a hydrodynamic minimization argument to fix the gauge freedom of a global rotation). This allows us to calculate the movement of an attached GNP and determine the observed local cross-section rotation relative to the laboratory coordinate system $\omega_{no twist}(\phi,s)$. **(B)** Map of $\omega_{no twist}(\phi,s)$ as function of beat-cycle phase $\phi$ and arc-length *s*. **(C)** Rotation angle $\omega_{no twist}(\phi)$ as function of beat-cycle phase for different arc-length positions *s* (color). From each curve, we obtain a peak-to-peak amplitude. **(D)** Peak-to-peak amplitudes ${\Delta\omega}_{no twist}(s)$ as function of arc-length *s*. The average over all arc-lengths equals 7.2° ± 0.3° (mean ± SEM, n = 28).

Goldstein, Raymond E, Thomas R Powers, and Chris H Wiggins. 1998. “Viscous Nonlinear Dynamics of Twist and Writhe.”

Happel, John, and Howard Brenner. 1983. 1 *Low Reynolds Number Hydrodynamics*. Dordrecht: Springer Netherlands. https://link.springer.com/10.1007/978-94-009-8352-6 (June 21, 2023).

Jikeli, Jan F. et al. 2015. “Sperm Navigation along Helical Paths in 3D Chemoattractant Landscapes.” *Nature Communications* 6(1): 1–10.

Mojiri, Soheil et al. 2021. “Rapid Multi-Plane Phase-Contrast Microscopy Reveals Torsional Dynamics in Flagellar Motion.” *Biomedical Optics Express* 12(6): 3169–80. http://opg.optica.org/boe/abstract.cfm?URI=boe-12-6-3169.

Ruhnow, Felix, David Zwicker, and Stefan Diez. 2011. “Tracking Single Particles and Elongated Filaments with Nanometer Precision.” *Biophysical Journal* 100(11): 2820–28.

Solovev, Anton, and Benjamin M. Friedrich. 2021. “Lagrangian Mechanics of Active Systems.” *European Physical Journal E* 44(4): 1–15.
